# Establishing the molecular basis for MDA5 mutation-linked autoimmunity

**DOI:** 10.64898/2026.01.20.700587

**Authors:** Ling Xu, Harim Jang, Kevin Chung, Rong Guo, Alisia Pan, Anna Marie Pyle

**Affiliations:** Department of Molecular, Cellular and Developmental Biology, Yale University, New Haven, CT 06511, USA; Howard Hughes Medical Institute, Chevy Chase, MD 20815, USA; Department of Chemistry, The Wertheim UF Scripps Institute, Jupiter, FL 33458, USA; Department of Molecular Biophysics and Biochemistry, Yale University, New Haven, CT 06511, USA; Department of Chemistry, Yale University, New Haven, CT 06511, USA

**Author notes:** Correspondence (L. X.) and (A. M. P.).

## Abstract

Melanoma differentiation-associated protein 5 (MDA5), a member of the RIG-I-like receptor family, is a cytoplasmic sensor essential for innate antiviral immunity. MDA5 distinguishes viral RNA from host RNA in part through its ATP hydrolysis activity, which promotes filament turnover on shorter endogenous dsRNAs. Here, we show that the gain-of-function T331I disease-linked mutation within the ATP binding pocket disrupts this balance, resulting in constitutive interferon signaling. Through a combination of cryo-electron microscopy (cryoEM), biochemical assays, and cellular analyses, we reveal the extensive network of interactions that precisely position ATP for catalysis in the wild-type MDA5 ATP binding pocket, and also demonstrate that the T331I mutation impairs ATPase activity, thereby stabilizing MDA5-dsRNA complexes and leading to aberrant immune activation. These findings elucidate how MDA5 ATPase activity regulates antiviral specificity and prevents autoimmunity by controlling filament stability and downstream signaling, offering a mechanistic molecular explanation for disease pathogenesis.

**GRAPHICAL ABSTRACT:** 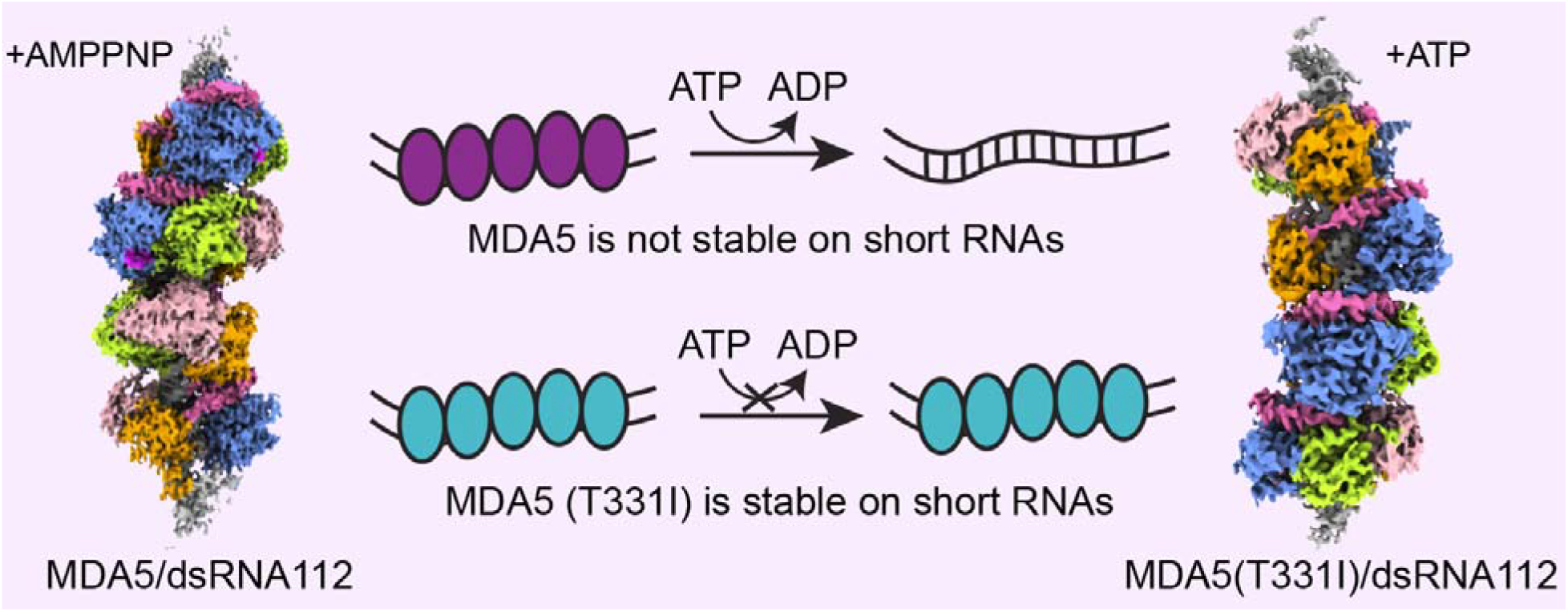

## INTRODUCTION

Melanoma differentiation-associated protein 5 (MDA5), encoded by the *IFIH1* gene, is a critical component of the vertebrate innate immune system, playing an important role in detecting and responding to viral infections. As a member of the RIG-I like receptor (RLR) family, MDA5 specifically recognizes long double-stranded RNA molecules (dsRNA), which are common features of many viral genomes and replication intermediates(1–3). Upon activation, MDA5 triggers a signaling cascade that culminates in the production of type I interferons (IFNs) and proinflammatory cytokines, generating a robust antiviral response(2,4). The importance of MDA5 in antiviral immunity is underscored by its involvement in defending against a broad range of viruses, including picornaviruses such as encephalomyocarditis virus (EMCV) and foot-and-mouth disease virus (FMDV)(2,4), and coronaviruses such as SARS-CoV2(5–8). MDA5 binds cooperatively to long dsRNA, ultimately forming stable filaments that interact with adapter protein MAVS to initiate downstream signaling(2). The length of MDA5-dsRNA filaments is regulated by MDA5-catalyzed ATP hydrolysis, which induces filament deterioration, thereby providing a proofreading mechanism that enables MDA5 to distinguish between long viral and short cellular dsRNAs(9–12). While the specific pathological link between MDA5 function and autoimmune diseases remain challenging to decipher given their wide spectrum of clinical presentations, recent studies have identified mutations in the *IFIH1* gene encoding MDA5 that is associated with interferon-driven autoinflammatory conditions(2,13-19). For example, the gain-of-function mutation c.2465G>A (p.Arg822Gln), in *IFIH1* was reported to cause Singleton-Merten syndrome. Additionally, Aicardi-Goutières syndromes are related to heterozygous missense variants in *IFIH1* (such as c.1465G>A, p.Ala489Thr)(20,21). These mutations lead to aberrant activation of MDA5, resulting in inappropriately high levels of interferon signaling. Of particular interest is a missense variant c.992C>T (p.Thr331Ile) in *IFIH1* that is associated with Singleton-Merten syndrome(22). The T331I mutation is linked to increased interferon production and autoimmune diseases(19,22). Previous studies suggest that T331I induces the IFN signaling pathway in the absence of exogenous dsRNA ligands and triggers a more robust interferon response than wild-type MDA5 in the presence of stimulating of poly (I:C) or dsRNAs as short as 162-bp(22).

While these results are notable, the actual molecular mechanism underlying constitutive MDA5 T331I signaling remains unclear. Thr331 is of particular interest because it is located in the highly conserved Motif I region of the MDA5 Hel1 domain, where precise amino acids are crucial for RNA-dependent ATP hydrolysis by Superfamily II (SF2) RNA-dependent ATPase enzymes(23). Therefore, the T331I mutant is likely to provide insights into the specific role of ATP hydrolysis in enabling MDA5 to maintain a balance between specific antiviral response and nonspecific autoimmune activation(9,24,25). It is thought that MDA5-catalyzed ATP hydrolysis facilitates rapid disassociation of MDA5 from relatively short endogenous dsRNAs while enabling it to remain bound to viral RNAs long enough to activate signaling(26). Despite the recognized importance of ATP hydrolysis in MDA5 function and filament stabilization, there is a gap in our understanding of the structural link between MDA5 catalytic function and its ability to form stable dsRNA complexes capable of signaling. We address this critical knowledge gap by systematically investigating the impact of the MDA5 T331I mutation on dysregulated signaling through comprehensive enzymatic, biochemical and cell-based studies coupled with high-resolution structural analysis of receptor-RNA complexes. Through this strategy, we establish the role of the ATPase active site in MDA5 discrimination between host and pathogen RNA molecules, and we directly visualize the short, functional T331I/RNA complexes that result from deactivation of MDA5 ATPase activity. These findings provide new insights into the basic mechanism of RNA-stimulated MDA5 signaling and the processes that dysregulate MDA5 behavior.

## MATERIAL AND METHODS

### Cloning, Expression and Purification of Human MDA5 Protein

Both full-length human MDA5 recombinants and MDA5 (ΔCARDs) were cloned into pET-SUMO expression vector using common molecular cloning techniques and verified by DNA sequencing. Site-specific mutations were introduced by Q5 Site Directed Mutagenesis Kit (NEB) using appropriate primers to make MDA5 T331I mutant construct using MDA5(ΔCARDs) plasmid. The full-length MDA5 was expressed in *E*.*coli* strain Rosetta (DE3) using LB medium at 16°C for 4 hours after adding 0.2 mM IPTG, while both MDA5 (ΔCARDs) and MDA5 T331I mutant were grown at 16°C for 16 hours after IPTG induction. Cells were harvested and resuspended in binding buffer (25mM Na_2_HPO_4_, 1M NaCl, 2mM ß-ME, 10% glycerol, pH 7.4) supplemented with protease inhibitor. For full-length MDA5, 0.2% IGEPAL was added into the binding buffer to reduce the formation of MDA5 aggregates. Cells were lysed using Microfluidizer and clarified by centrifugation. Recombinant protein was then purified by nickel-chelating columns (GE Healthcare) and washed and eluted using 20 mM and 300 mM imidazole respectively in binding buffer. SUMO protease was added to the eluted fraction to remove the SUMO tag. Protein was further purified by ion exchange with HiTrap Heparin HP column (GE Healthcare) and by size exclusion with HiLoad Superdex 200 (16/600) columns (GE Healthcare). Protein was pooled and concentrated in storage buffer (25 mM HEPES pH 7.4, 200 mM NaCl, 10% glycerol, 5 mM ß-ME) before flash freezing and storage until use at −80°C.

### RNA Transcription and Purification

The DNA templates for transcribing double stranded RNA (dsRNA, sequence see in supplementary Table S2) were synthesized by IDT (Integrated DNA Technologies) and dissolved in ultrapure water to a final concentration of 60 mM. *In vitro* transcription of the dsRNA was performed using in-house-prepared T7 RNA polymerase in a transcription buffer containing 40 mM Tris-HCl pH 8.0, 10 mM NaCl, 23 mM MgCl2, 2 mM spermidine, 0.01% Triton X-100, 10 mM DTT and 5 mM each NTP. 0.3uM DNA template was added for each 1 mL of transcription and the reaction was incubated at 37°C for 2 hours. The transcribed dsRNAs were then mixed with 2x urea loading dye containing bromophenol blue and xylene cyanol, which was then purified on 10% denaturing polyacrylamide gels (19:1 acrylamide/bisacrylamide). The band was visualized with UV shadowing, cut with a sterile blade, crushed with a sterile syringe and eluted in a gel elution buffer (10 mM Na-MOPS, pH 6.0, 300 mM NaCl and 1 mM EDTA) overnight. The eluted RNA was then ethanol precipitated, resuspended in an RNA storage buffer (6 mM Na-MES pH 6.0) to a final concentration of 100 μM and frozen at −80°C for preparation of RNP samples. Before use in assays, RNA was diluted and heated to 95 °C for 2 min and cooled at 25 °C for 10 min to properly fold.

### Luciferase Activity Assay

Human pUNO-hMDA5 vectors containing full-length wild-type MDA5 for expression in mammalian cell culture were purchased from Invivogen. Mutations were introduced by Q5 Site Directed Mutagenesis Kit (NEB) using appropriate primers. HEK293T cells were grown in Dulbecco’s Modified Eagle Medium (Thermo Fisher Scientific) supplemented with 10% heat-inactivated Bovine Calf serum (Hyclone) and Non-Essential Amino Acids (Thermo Fisher Scientific). For IFN-β induction assays, 500,000 cells were seeded in each well in 24-well assay plates. After 24 h, each well was transfected with various concentrations of wild type or mutant expression pUNO-hMDA5 plasmids, 6 ng pRL-TK renilla luciferase reporter plasmid (Promega) and 150 ng of an IFN-β/Firefly luciferase reporter plasmid using Lipofectamine 2000 transfection reagent (Thermo Fisher Scientific). Cells were incubated for 24h before transfection with 1 µg/well of various RNAs. After 3h, 6h, 12 h or 24h, growth medium was aspirated and 100 uL of 1x passive lysis buffer (Promega) was added and incubated for 15 min at room temperature. Dual-Luciferase Reporter Assay System (Promega) and Biotek Synergy H1 plate reader (Biotek) were used to read luciferase measurements.

### RT-qPCR

Cells were prepared under the same transfection conditions as described in the Luciferase Activity Assay. Total RNA was extracted from cells using TRIzol reagent (Invitrogen) according to the manufacturer’s instructions. RNA samples were treated with DNase I (NEB) according to the manufacturer’s instructions, followed by an additional extraction with Acid-Phenol, pH 4.5 (Invitrogen). RNA concentration was measured using a NanoDrop spectrophotometer (Thermo Fisher Scientific), and 2 μg of total RNA was used for reverse transcription with SuperScript II reverse transcriptase (Thermo Fisher Scientific). The resulting cDNA was diluted and used for qPCR with LightCycler 480 SYBR Green I Master (Roche) on a CFX Opus 384 Real-Time PCR System (Bio-Rad). Transcript levels were normalized to endogenous ACTB and expressed relative to the condition without ectopic MDA5 expression and without RNA stimulation, which was set to 1.

### Western Blot

Cells were prepared under the same transfection conditions as described in the Luciferase Activity Assay. Western blot of wild-type MDA5 and T331I mutant used samples from the luciferase assay. Expression levels of MDA5, T331I and GAPDH were analyzed by immunoblotting with specific antibodies. Mouse anti-MDA5 antibody was used to detect the expression of MDA5. GAPDH was used to act as control.

### Electrophoretic Mobility Shift Assay (EMSA)

0.5 μM purified MDA5 and T331I mutant proteins were incubated with 20 nM 112bp dsRNA at room temperature for 30 min in binding buffer containing 25 mM HEPES (pH 7.5), 150 mM NaCl, 1.5 mM MgCl_2_. ATP was then added to final concentrations of 1 mM, 3 mM, or 5 mM, followed by an additional 6-minute incubation at room temperature. Subsequently, 2 μL loading buffer (80% glycerol) was mixed with each 10 μL sample, and the mixtures were loaded onto an SDS-PAGE gel and stained with GelRed. Gels were imaged by detecting GelRed using Bio-Rad ChemiDoc Touch Imaging System.

### ATPase Activity Assay

The ATPase activities of human wild-type MDA5 and T331I mutant were measured in the presence of poly (I:C) (Invitrogen) or invitro transcribed SLR50. Reactions were performed at 25°C using a standard method as previously described(27). In the pyruvate kinase-lactase dehydrogenase assay, hydrolysis of ATP is coupled to the oxidation of NADH, which is detected by measuring the decrease of absorbance at 340 nm. The reaction system contained 0.5 µM protein, 400 mM phosphoenolpyruvate, 10 U/ml pyruvate kinase, 10 U/ml lactate dehydrogenase, 2 mM ATP, 2 mM MgCl2, 0.25 mM NADH and 1 µM RNA in reaction buffer (10 mM Hepes, 75 mM NaCl, pH 7.5). The reaction was monitored using a Biotek Synergy H1 plate reader. Data from three independent experiments were analyzed using GraphPad Prism.

### Complexes Assembly for EM

The N-terminal 297 residues were removed from both protein constructs to reconstitute wild-type MDA5/dsRNA112 or T331I/dsRNA112 complexes. The protein-dsRNA complexes were first assembled by incubation 5uM proteins and 0.2uM dsRNAs at 4°C overnight in 25□mM HEPES, pH 7.4, 15025□mM NaCl, 2mM DTT, 2mM AMPPNP or ATP, 5mM MgCl_2_. The short filament complexes formed by protein and dsRNAs were verified by negative staining and then analyzed by Cryo-EM.

### Negative Stain Analysis

The wild-type MDA5/dsRNA112 or T331I/dsRNA112 complexes were applied to a glow discharged holey carbon grid that was precoated with a thin layer of continuous carbon film over the holes. After 45 s, the sample was wicked with filter paper and the grid was stained with three droplets of 2% (w/v) uranyl acetate solution. After another 45 s, the residual stain was blotted off, and the grid was air-dried. The EM data for the negatively stained specimen was collected on a FEI Talos L120C transmission electron microscope operated at 120-kV. Images were collected at a nominal magnification of 75,000×, with a total dose of approximately 20 e-/Å2 and defocus value ranging from −1.5 to −1.6 μm on a 4k×4k CetaTM 16M camera, with a pixel size of 1.97 Å on the object scale.

### Cryo-grid Preparation and Data Collection

For cryo-EM analysis of the wild-type MDA5/dsRNA112/AMPPNP and T331I/dsRNA112/ATP complexes. 4 μL sample was loaded onto glow discharged (60s, 25mA) QuantiFoil UtraAu R2/2 200-mesh grids. The grids were blotted for 2.5s in 100% humidity at 22°C with a blotting force offset of −4 and rapidly frozen in liquid ethane using Vitrobot (Thermo Fisher). Grids were screened with Glacios microscope operated at 200 keV and grids with good particle density and ice thickness were selected for data acquisition.

For cryoEM data collection, frozen grids were loaded into a Titan Krios microscope (FEI) operated at 300 kV, equipped with a post-GIF K3 summit direct detection camera. For MDA5/dsRNA112/AMPPNP complex, 5640 micrographs were collected. A defocus range of −0.5 µm and −2.0 µm were used. Each micrograph was collected at 30 frames per second with a total exposure time of 1.4s and a frame exposure time of 0.04s, resulting in a total dose of 51.98 e-/A2. For T331I/dsRNA112/ATP complex, 5990 micrographs were collected. A defocus range of −0.5 µm and −2.0 µm were used. Each micrograph was collected at 30 frames per second with a total exposure time of 1.5s and a frame exposure time of 0.03s, resulting in a total dose of 50.49 e-/A2.

### CryoEM Data Processing

Recorded movie frames were processed using cryoSPARC v4.0. Motion correction and CTF estimations were performed using default parameters in cryoSPARC. Exposures were curated and micrographs with obvious ice contamination, large motions, or damaged areas were removed. Particle picking was done with the automated blob picker and filtered using consecutive rounds of 2D classification. Particles were selected for initial model creation, generating three reconstructions, of which one was selected as the reference for further classification. In the first round of 3D classification, four classes were separated out. The target classes were subjected to further refinement. The CryoEM maps of filaments containing four MDA5 subunits bound to dsRNA were solved to resolution of 3.04 A□ for TetraMDA5/dsRNA and 3.39 AC for TetraT331I/dsRNA. For reconstruction, separate masked local refinements, and local and global CTF refinement were conducted on one subunit or two subunits to improve resolution of each. Two different maps were both obtained for the wild-type MDA5/dsRNA112/AMPPNP and T331I/dsRNA112/AMPPNP mutant. With the reconstructions which have resolutions of 2.96 AC and 2.81AC for wild-type DiMDA5/dsRNA and MonoMDA5/dsRNA structures, and 3.21 A□ and 3.12A□for DiT331I/dsRNA and MonoT331I/dsRNA complex, respectively (Fig. S1 and S8, Table S1).

### Model Building and Refinement

Model building was initiated by docking AlphaFold(28,29) predicted human MDA5 protein model into the generated reconstructions using UCSF Chimera(30). The models were then manually rebuilt in COOT to accommodate for the changes in the sequence of amino acids. The final models were improved by iterative rounds of real-space refinement against the sharpened cryoEM map in PHENIX using Ramachandran and rotamer restraints for protein chains, and subsequent rebuilding in COOT(30–32). Model building and validation statistics are listed in Table S1. The model were then analyzed using UCSF ChimeraX(33).

## RESULTS

### MDA5 disease mutant T331I can be stimulated by both short and long dsRNAs

Previously studied pathogenic MDA5 variants have been shown to enhance the stability of MDA5-dsRNA filaments either by diminishing ATPase activity or increasing RNA binding affinity, resulting in enhanced interferon signaling via interactions with host RNAs(13,26). To determine the functional consequences of the MDA5 disease mutant T331I, we monitored its behavior in well-established cell-based reporter systems for interferon signaling(34) under diverse conditions, and in the presence of poly(I:C) or diverse dsRNAs (see methods and supplementary information) (Fig. 1A-1C and Fig. S2). A comparative analysis of wild-type MDA5 and the T331I mutant yielded interesting insights. For example, under various concentrations of poly (I:C) after a 16-hour induction, we observed that the T331I mutant consistently showed an approximately 2-fold increase in luciferase activity relative to wild-type MDA5 (Fig. 1A and Fig. S3A). Notably, in the absence of poly (I:C) stimulation, the T331I mutant still displayed a high level of interferon induction (Fig. 1A). These findings indicate that the T331I mutation can stimulate interferon signaling even in the absence of exogenous RNA and that it has enhanced overall signaling capability. This is consistent with previous research indicating that different substitutions of the threonine residue at position 331 are consistently linked to increased IFN-β expression in patients(22).

**Figure 1.**
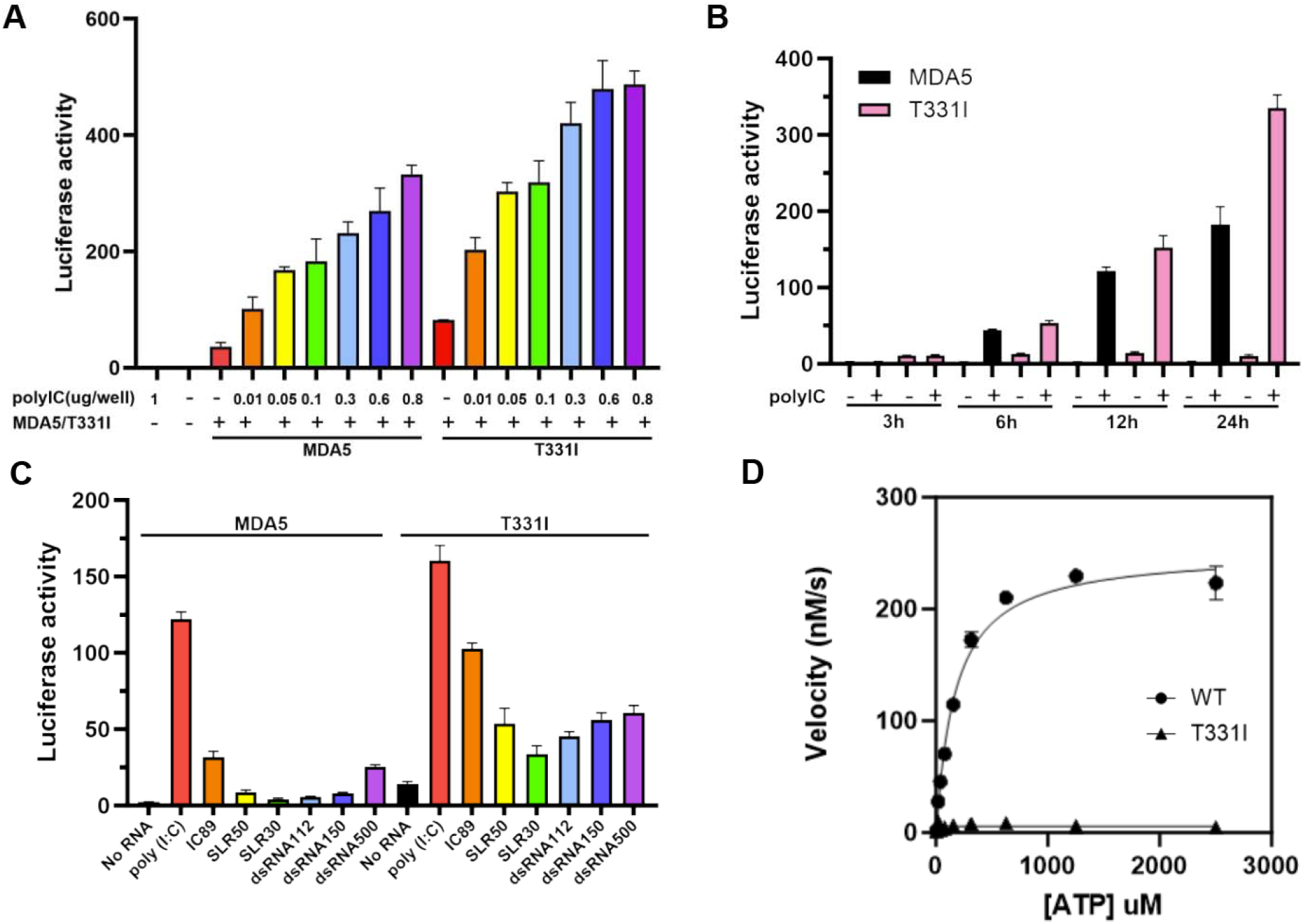
ATPase-deficient MDA5 T331I mutant exhibits hyperactivity in cell-based IFN-β reporter assays. A. IFN-β induction by MDA5 and the MDA5 T331I mutant upon poly(I:C) stimulation in HEK293T cells. Cells were transfected with increasing amounts of poly(I:C) (0.01, 0.05, 0.1, 0.3, 0.6, and 0.8 µg). Luciferase activities were measured 12 h post-transfection. Data are represented as mean +/− SD (n=3 biological replicates). Data were analyzed by ordinary one-way ANOVA for multiple comparisons in Prism. B. Time course of IFN-β induction by MDA5 and MDA5 T331I mutant stimulated with 1µg/well poly (I:C) in HEK293T cells. Luciferase activities were measured at 3, 6, 12, and 24 hours post-transfection for both MDA5 and T331I mutant. C. IFN-β induction by MDA5 and MDA5 T331I mutant stimulated with the indicated RNAs in HEK293T cells. 1 ug/well RNAs were transfected. RNA transfected were poly (I:C), IC89 which is an 89 bp *in vitro* transcribed double-stranded RNA composed exclusively of inosine (I) and cytosine (C) nucleotides (IC89). 50 bp and 30 bp stem loop RNA (SLR50 and SLR30, see sequences in Table S2) and double-stranded RNA molecules with blunt ends, comprising 112, 150, or 500 nucleotides per strand (dsRNA112, dsRNA150 and dsRNA500). D. ATPase activity of wild-type MDA5 and T331I mutant induced with poly (I:C). The assay was performed in triplicate.

To explore potential kinetic differences in the signaling behavior of wild-type MDA5 and T331I, we compared their temporal dynamics of interferon induction. Cells with wild-type MDA5 or T331I mutant plasmids were stimulated with poly (I:C) for 3, 6, 12 and 24 hours (see Methods). At the early time point of three hours, neither wild-type MDA5 nor the T331I mutant showed a significant IFN response (Fig. 1B), suggesting that the activation of MDA5 is slower than the related RIG-I receptor, at least when stimulated by transfected RNA(35). At both 6-hour and 12-hour time points, interferon induction by T331I was slightly higher than that of wild-type MDA5. 24 hours after poly(I:C) stimulation, a 2-fold increase was observed in the interferon induction by T331I relative to the wild-type MDA5 (Fig. 1B). Importantly, the basal signal level of T331I without the addition of exogenous RNA remained steady from 3 hours to 24 hours, suggesting that activation of the T331I mutant by the RNA might be regulated by a different mechanism.

To evaluate the importance of dsRNA length in activating MDA5 signaling and to determine how this is affected by the T331I mutation, we measured interferon induction by wild-type MDA5 and T331I cell-based reporter systems using RNAs of increasing length. Previous studies have established that MDA5 binds cooperatively to long dsRNAs in a sequence-independent manner, forming extended filaments that contribute to a downstream antiviral signaling pathway(24,36-38). Here, we showed that MDA5 exhibits high levels of RNA-stimulated interferon induction upon transfection with low-molecular weight poly (I:C) (average length 0.2-1kb), and moderate levels upon transfection with a short designed polyIC duplex (IC89) or a long random-sequence dsRNA that is 500 bp in length while shorter dsRNAs showed negligible activity (Fig. 1C). By contrast, the T331I mutant exhibited robust signaling upon stimulation by almost every RNA tested, including dsRNAs as short as 30 base pairs (SLR30) in length (Fig. 1C). This indiscriminate activity of T331I implies that the Threonine to Isoleucine mutation enables MDA5 to form signaling-competent filaments on dsRNAs of almost any length or sequence, having completely lost its ability to discriminate against short host transcripts. Given that the site size of MDA5 on dsRNA is about 13-14 base pairs, oligomers as short as two T331I units were sufficient for signaling. This indicates that the T331I mutant formed exceptionally stable complexes from which MDA5 cannot readily dissociate, thereby allowing even very short T331I-dsRNA complexes to become stabilized and engage in the signaling pathway.

### MDA5 T331I mutant lacks ATPase activity, and its RNA filaments cannot disassemble

It is well established that filaments formed by MDA5 on long double-stranded RNA undergo disassembly upon MDA5-catalyzed ATP hydrolysis(9). This process is believed to tune the stability of the MDA5 filament in a length-dependent manner, enabling stable complex formation on very long viral dsRNAs while preventing the formation of stable MDA5 complexes with the abundant short dsRNAs in the cytoplasm of host cells. Previous studies have also indicated that ATPase-deficient MDA5 mutants can increase the affinity of MDA5 for short dsRNAs(13,20). Despite the compelling mechanistic model suggested by these reports, stable mutant filaments have never been visualized nor proven to be a specific outcome of disruption in ATP binding pocket of MDA5.

To investigate the biochemical consequences of the MDA5 T331I mutation, we purified both wild-type MDA5 and T331I mutant proteins and examined their ATPase and filament-forming activities. As in previous studies, poly (I:C) was employed as the activating dsRNA ligand(37). Consistent with reported observations, wild-type MDA5 exhibited robust ATPase activity upon binding to poly (I:C). However, the T331I mutant displayed no detectable ATPase activity in the presence of poly (I:C) (Fig. 1D). We therefore examined whether constitutive activity of the T331I mutant results from impaired filament disassembly, even on short RNAs.

To investigate the nature of T331I filaments and to compare them with filaments formed by wild-type MDA5, we employed negative-staining and cryoEM. Both wild-type MDA5 and T331I form individual protein monomer particles in the absence of dsRNA (Fig. 2A and Fig. S3B). Importantly, both wild-type MDA5 and T331I mutant formed well-ordered filaments on a relatively short 112bp dsRNA (dsRNA112) in the presence of AMPPNP without displaying nonspecific aggregation, suggesting that both proteins form stable, uniform complexes on dsRNA in presence of AMPPNP (Fig. 2B). Upon replacing AMPPNP with ATP, an abundance of free RNA molecules were observed in both cryoEM and EMSA data, and only a few filaments were formed by wild-type MDA5, suggesting that MDA5-catalyzed ATP hydrolysis triggered rapid disassembly of wild-type MDA5 filaments on dsRNA112 (Fig. 2C, top panel, and Fig. S4A). In contrast, the short filaments formed by T331I on dsRNA112 were stable even in the presence of ATP and no evidence of filament disassembly was observed (Fig. 2C, bottom panel, and Fig. S4B). Based on these results, we conclude that the T331I mutation eliminates the ATPase activity of MDA5, which in turn blocks the disassembly of filaments formed by the T331I mutant even on short dsRNA. This stabilization of small filaments explains the hyperresponsiveness of T331I to endogenous short RNA ligands and the prominent signaling in the presence of relatively short RNAs in our cell-based reporter assay (e.g., IC89, SLR50, dsRNA112 and dsRNA150; Fig. 1C).

**Figure 2.**
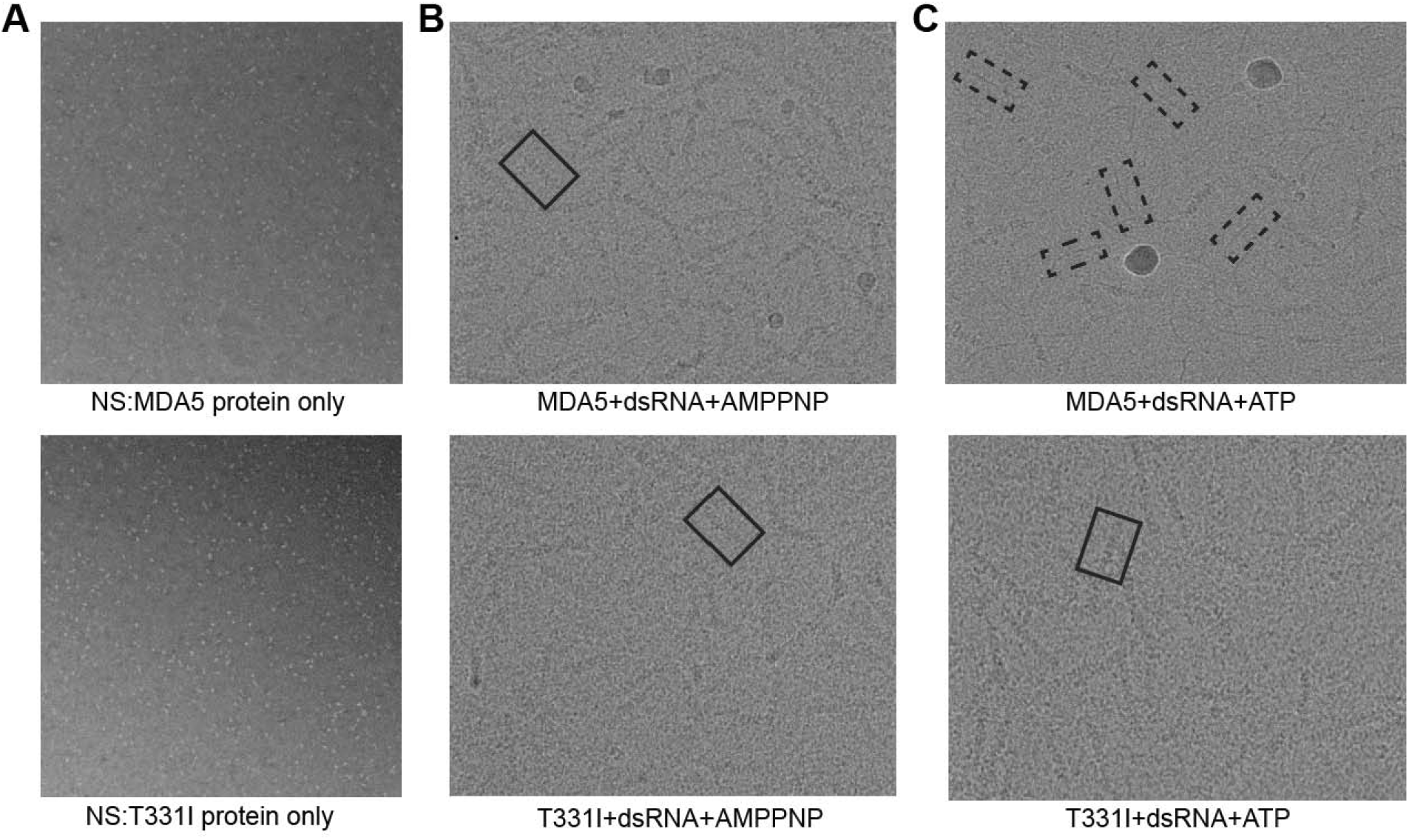
Impaired ATP-dependent disassembly of MDA5 and T331I/RNA filaments. A. Negative-stain electron micrographs of wild-type MDA5 (top) and the T331I mutant (bottom) in the absence of dsRNA. Small white dots indicate the individual protein alone particles on the grid. B. CryoEM micrographs showing filaments on the RNA formed by wild-type MDA5 (top) and the T331I mutant (bottom) assembled on dsRNA112 in the presence of the ATP analogue AMPPNP. Boxes highlight representative repetitive, punctate patterns of protein domains in linear filaments formed by MDA5/dsRNA112 (top) and T331I/dsRNA112 (bottom). Images are representative of at least three independent micrographs. C. CryoEM micrographs of wild-type MDA5 (top) and the T331I mutant (bottom) with dsRNA112 in the presence of ATP. Dashed-squares indicate free dsRNA molecules. Solid squares indicate dsRNA-protein filaments formed by T331I/dsRNA112 in presence of ATP.

### CryoEM elucidation of MDA5 bound to a short dsRNA

To further investigate the mechanism by which short RNA filaments stimulate the innate immune response, we examined both wild-type MDA5 and T331I mutant filaments using cryoEM. We first visualized the domain architecture and nucleotide binding pockets of wild-type MDA5 in complex with short dsRNA using cryoEM. To minimize nonspecific protein precipitation, we removed the N-terminal Caspase Recruitment Domains (CARDs) from the MDA5 constructs. Previous studies(38), including our own, have demonstrated that MDA5 constructs lacking CARDs retain the ability to form filaments on dsRNA (Fig. 2B). We then assembled complexes between wild-type MDA5 and a 112 bp dsRNA, loaded the samples onto cryoEM grids, and subjected them to our imaging pipeline (see Fig. S1 and Methods). To address heterogeneity in filament length, particles from both samples were classified into several groups during 3D reconstruction (Fig. S1). We selected filaments containing at least four protein subunits (TetraMDA5/dsRNA complexes) for further refinement (Fig. 3A). This approach provided high-resolution structural information for specific regions of the filament, especially the ATP-binding pocket and the MDA5 monomer-monomer interface. The refinements were focused either on the central subunit (MonoMDA5/dsRNA complex) or on two adjacent subunits (DiMDA5/dsRNA complex) within the TetraMDA5/dsRNA complex (Fig.□3B and 3C).

**Figure 3.**
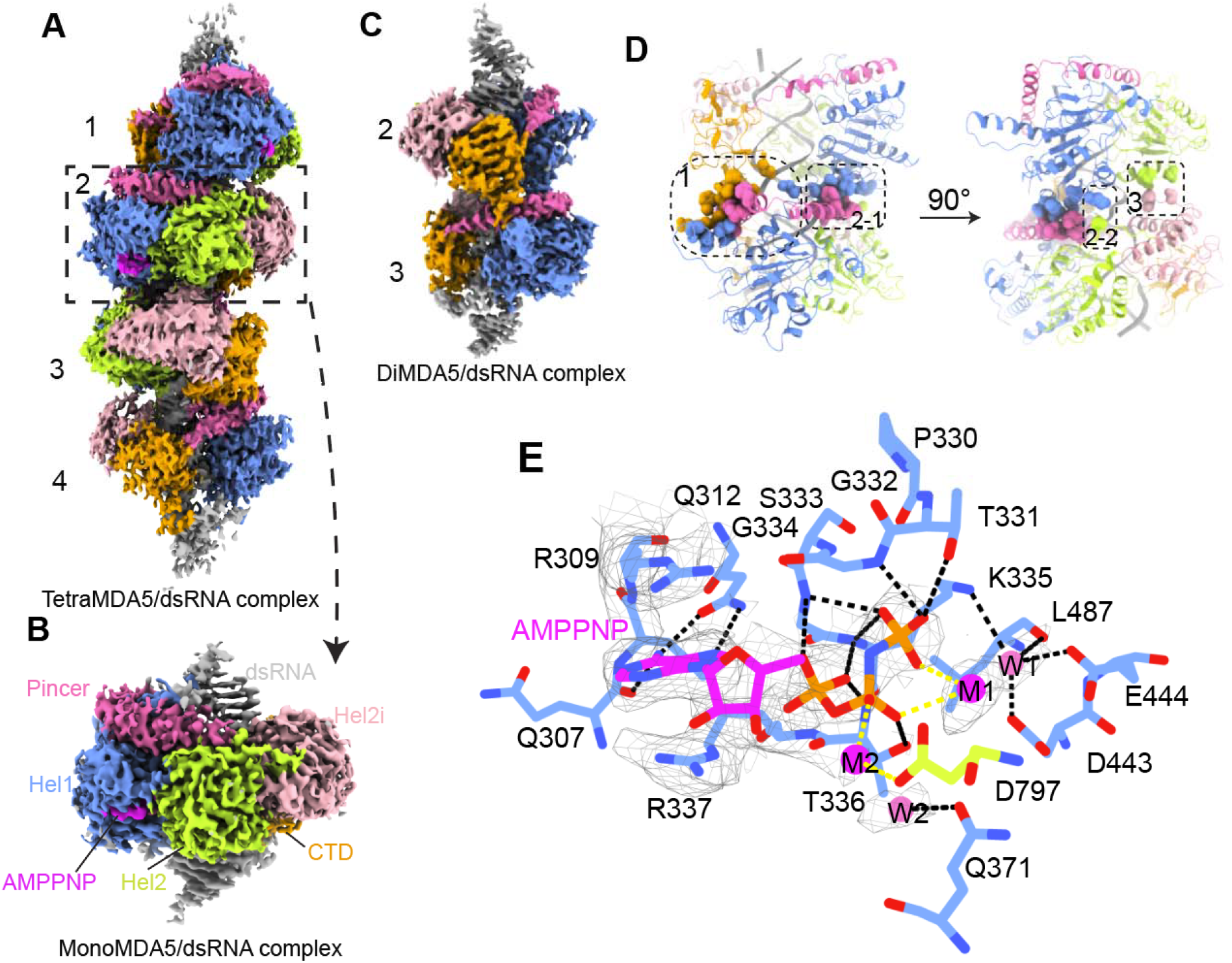
CryoEM structure of wild-type MDA5/dsRNA complex in the presence of AMPPNP. A. CryoEM map of a short filament formed by wild-type MDA5 and short dsRNA112. B. High-resolution structure of a single wild-type MDA5 molecule bound to dsRNA in the presence of AMPPNP. Domain coloring: Hel1 (pale blue), Hel2 (limon), Hel2i (pink), pincer domain (magenta), C-terminal domain (orange), and RNA (gray). C. Refined cryo-EM map showing two wild-type MDA5 subunits bound to dsRNA. D. Atomic model corresponding to the structure depicted in panel C. The monomer-monomer interfaces are indicated with dashed boxes. Details of the four interaction clusters can be found in Fig. S5. E. AMPPNP binding pocket (magenta) with two magnesium ions as magenta spheres and two water molecules as pink spheres. At the current resolution, a plausible M2 ion is assigned to the density adjacent to the AMPPNP-bound M1. Interaction network within the nucleotide binding pocket of wild-type MDA5. Thr331 forms a polar contact with the γ-phosphate.

We observed that wild-type MDA5, in the presence of AMPPNP, forms closed rings around short dsRNA (Fig. 3A). The twist of this short filament resembles that of MDA5 filaments assembled on long dsRNAs reported previously(38). The TetraMDA5/dsRNA and DiMDA5/dsRNA complex structures achieved overall and local resolutions of approximately 3.0 Å and 2.96 Å, respectively, while the MonoMDA5/dsRNA complex reached an overall and local resolution of approximately 2.81 Å (Table S1). These high-quality maps enabled us to construct accurate atomic models for both the DiMDA5/dsRNA and MonoMDA5/dsRNA complexes (Fig. 3B and 3C). The high-resolution DiMDA5/dsRNA model allowed us to investigate the monomer-monomer interface between two MDA5 subunits within the filament (Fig. S2A to S2C). In the presence of AMPPNP, the two adjacent MDA5 subunits twist about 82.8° on the dsRNA, forming three distinct clusters of protein-protein interactions (Fig. 3D). The pincer domain acts as an anchor, contributing to the monomer-monomer interface, as previously described in mouse MDA5 long filaments(38). Notably, the human MDA5 monomer-monomer interface involves a greater number of amino acids at these anchor points than that in mouse MDA5 filament(38) (Fig. S5). The high-resolution DiMDA5/dsRNA structure further revealed that the CTD tail of subunit 2 is fully engaged in forming the interface, interacting with both the pincer and Hel1 domains of subunit 3. Additionally, we identified a third interaction cluster formed by Hel2 of subunit 2 and Hel2i of subunit 3, which has not been previously reported (Fig. S5D). Mutagenesis analysis of amino acids on this surface revealed that substitution of Ser568 and Met570 with alanine reduces MDA5 luciferase activity (Fig. S6), indicating that these residues contribute to intersubunit affinity. Together, three major clusters of interactions create a monomer-monomer interface area of approximately 840 Å^2^, as calculated by PDBePISA(39), and these help to maintain the helical twist of MDA5 during filament assembly on dsRNA.

Furthermore, the high-resolution structure of the MonoMDA5/dsRNA complex enabled us to visualize the nucleotide binding pocket in remarkable detail, revealing that numerous amino acids contribute to its formation (Fig. 3I). This critical pocket contains several hydrophilic residues that constitute motifs Q and I (Fig. S7), characteristic features of DExD/H-box RNA helicases in superfamily 2(23). Within the MonoMDA5/dsRNA complex, AMPPNP is nestled in a cleft formed by residues from the Hel1 domain, with these interactions precisely positioning the nucleotide. Arg309 and Arg337 create a classic sandwich interaction with the adenine base of AMPPNP, while Glu307 and Glu312 form polar contacts that further stabilize the adenine. Additionally, the P-loop, formed by residues Thr331 to Gly334, stabilizes the phosphate tail of AMPPNP.

Two magnesium ions (M1 and M2) are clearly resolved: M1 coordinates with both the β- and γ-phosphates of AMPPNP, as previously observed in SF2 helicases(40) (Fig. 3E), while M2 coordinates with the β-phosphate and Asp797 from Hel2. Although water molecules close to the γ-phosphate could not be resolved at this resolution, we report for the first time that both Asp443 and Glu444 of the conserved DECH motif coordinating a water molecule (W1) in the pocket. This water also forms polar contacts with the side chain of Lys335 and the backbone of Leu487. A second water molecule (W2) was identified near Gln371. Together, these interactions define the nucleotide binding pocket and precisely position AMPPNP within the structure.

### Stable RNA-T331I filament forms on short dsRNA

We next sought to determine whether there might be a structural basis for the unusually active signaling behavior of T331I. Sequence analysis revealed that Thr331 is located within the nucleotide binding pocket of MDA5 and is part of Motif I, a region highly conserved among RNA helicases (Fig. S7). To visualize the effect of the threonine-to-isoleucine mutation on MDA5 filament formation, we solved the structure of the T331I mutant bound to dsRNA in the presence of ATP, as this mutant lacks ATPase activity. The construct was identical to wild-type MDA5, except for the threonine substitution by isoleucine at position 331. Data processing followed the same workflow as for the wild-type MDA5/dsRNA complex, including refinement of maps containing four MDA5 subunits bound to dsRNA (TetraT331I/dsRNA complex), which achieved overall and local resolutions of approximately 3.4 Å. Additionally, we refined maps with two (DiT331I/dsRNA complex) and one (MonoT331I/dsRNA complex) MDA5 subunits bound to dsRNA, reaching overall and local resolutions of about 3.2 Å and 3.12 Å, respectively (Fig. S8).

We observed that the T331I mutant formed short filaments on double-stranded RNA, similar to those seen in the TetraMDA5/dsRNA complex (Fig. 4A). High-resolution analysis of the DiT331I/dsRNA complex revealed three clusters of interactions between the two protein subunits, consistent with the wild-type MDA5/dsRNA complex (Fig. 4B). However, the monomer-monomer interface area between the two subunits is approximately 1264 Å^2^ in the DiT331I/dsRNA complex, which is about 424 Å^2^ larger than that of the DiMDA5/dsRNA complex, suggesting increased stability of the T331I/dsRNA filament in the presence of ATP.

**Figure 4.**
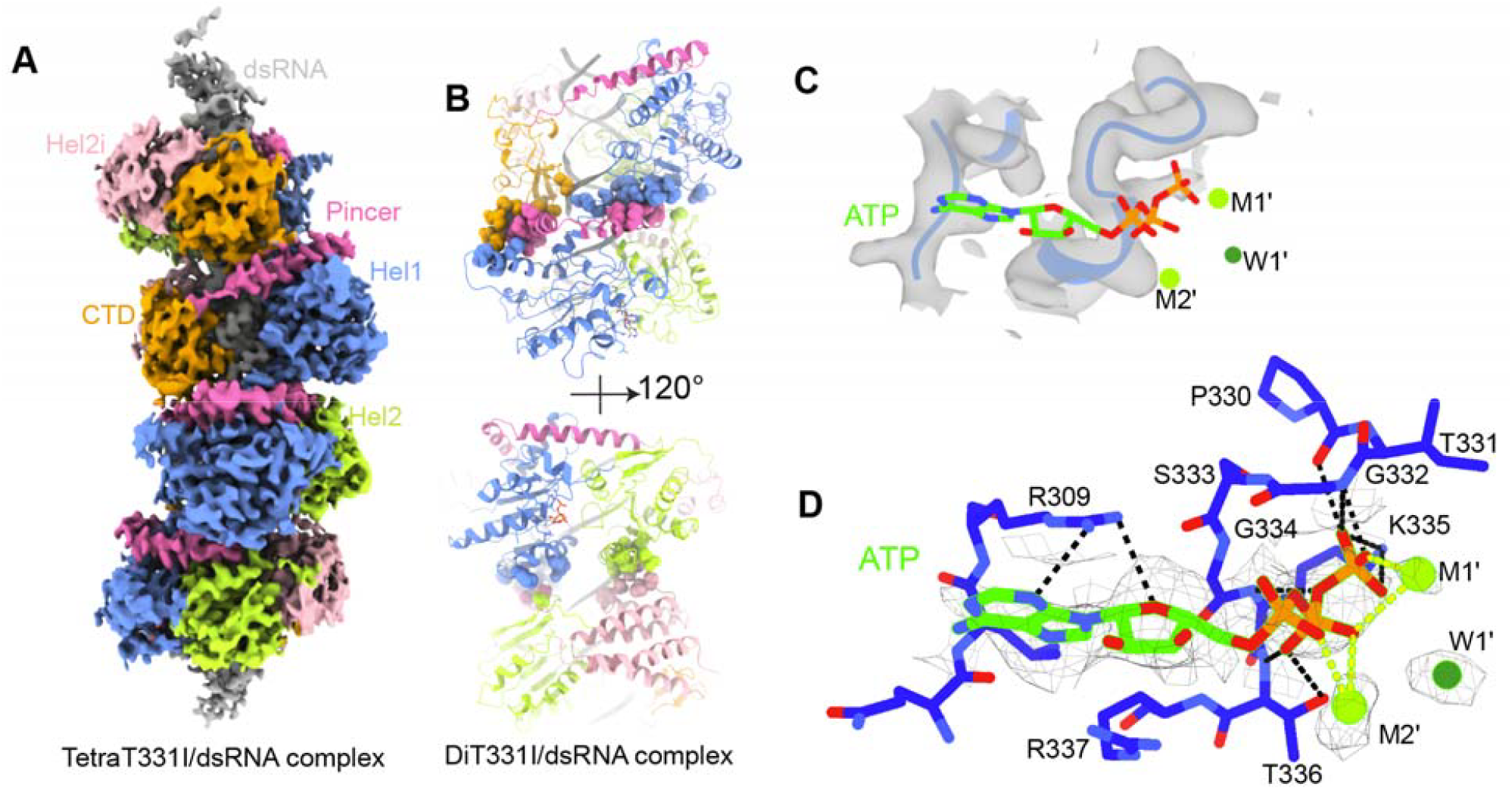
Structural analysis of ATP recognition by the T331I mutant. A. CryoEM map of a short filament formed by the T331I mutant and dsRNA112 in presence of ATP. B. Structural model showing two T331I subunits bound to dsRNA, with the monomer-monomer interfaces between protein units depicted as spheres. C. ATP molecule and its binding pocket in the T331I mutant. The density map highlights amino acid residues involved in ATP binding. D. Structural details of the interaction network among ATP, protein, and magnesium ions. A plausible M2’ ion is assigned to the density adjacent to the ATP-bound M1’.

Examining the ATP binding pocket in the high-resolution MonoT331I/dsRNA complex, we found that ATP occupies the same pocket as in the wild-type MonoMDA5/dsRNA complex, but it engaged with significantly fewer amino acid residues (Fig. 3E, 4C and 4D). For example, whereas the wild-type forms a classic sandwich interaction (Arg309-adenine base-Arg337), in the T331I mutant this stacking geometry is disrupted due to an outward shift of the ATP, resulting in altered positioning of Arg337 and leaving Arg309 as the primary residue maintaining polar contacts with the adenine base and sugar ring (Fig. 3E and 4D). Additionally, only three residues from the P-loop now interact with the phosphate tail, and Ile331 no longer forms a polar contact with the γ-phosphate of ATP (Fig. 3E and 4D), indicating increased flexibility in the ATP binding site. Furthermore, we were able to resolve two magnesium ions (M1’ and M2’): M1’ coordinates with both the β- and γ-phosphates of ATP, while M2’ coordinates with the α- and β-phosphates, suggesting strong metal-phosphate coordination. However, the single water molecule identified in the mutant structure is no longer associated with Asp443 and Glu444 of the DECH motif (Fig. 3E and 4D), indicating a potential impairment in ATP hydrolysis.

### T331I mutant forms a rigid filament

To determine whether the observed monomer-monomer interface difference between subunit 2 and subunit 3 in both wild-type DiMDA5/dsRNA and DiT331I/dsRNA extends to other subunits, we analyzed the helical twist values throughout each filament with four subunits. Although the subunits at both ends of the filament were of lower resolution, we successfully assigned the amino acids in the pincer domain for four subunits in the filament structure, allowing us to accurately define and compare helical twist values for the two MDA5 constructs (Fig. 5A and 5B). Interestingly, in the TetraMDA5/dsRNA complex, the helical twists between adjacent subunits varied, with values of approximately 79.94°, 82.30°, and 79.79° from subunit 1 to subunit 4, respectively (Fig. 5A). In contrast, the TetraT331I/dsRNA complex exhibited nearly identical helical twists between subunits in presence of ATP or AMPPNP, with measurements of 71.66° (subunit 1-2), 71.44° (subunit 2-3), and 71.51° (subunit 3-4) (Fig. 5B, Fig. S9 and Fig. S10B).

**Figure 5.**
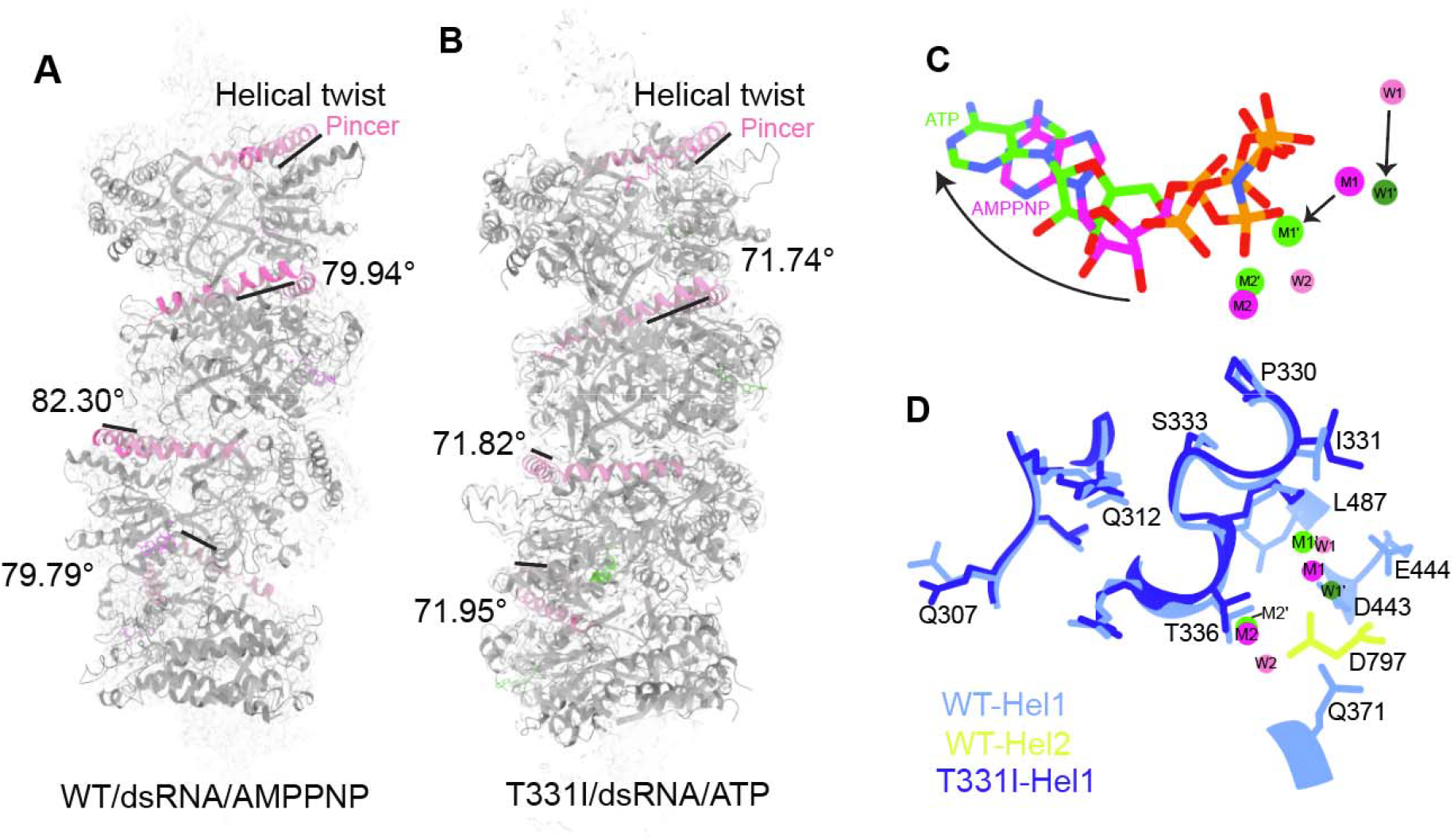
Structural differences between wild-type MDA5 and the T331I mutant. A. Wild-type MDA5 forms a filament with variable helical twist on short dsRNA. Although the overall resolution is insufficient to build a high-resolution model for all four subunits, the resolution is adequate to model the Pincer domain of each subunit. Helical twists were then readily measured using amino acids 869–882 of the pincer domain from each subunit. The helical twist angles between subunits are as follows: 79.94° (subunit 1 to 2), 82.30° (subunit 2 to 3), and 79.79° (subunit 3 to 4). B. The T331I mutant forms filaments with a constant helical twist on short dsRNA. The measured helical twist angles between subunits are 71.74° (subunit 1 to 2), 71.82° (subunit 2 to 3), and 71.95° (subunit 3 to 4). C. Overlay of nucleotide-binding sites: AMPPNP from the wild-type MDA5/dsRNA structure is compared with ATP from the T331I/dsRNA structure. Magnesium ions are indicated as M1 and M2 (colored in magenta) in the wild-type structure, and M1’ and M2’ (colored in green) in the T331I mutant. Water molecules are labeled W1 and W2 (colored in pink) in the wild-type, and W1’ (colored in dark green) in the mutant. D. Comparison of amino acids involved in nucleotide binding pocket and the conformational change in the wild-type MDA5 (amino acids in Hel1 are colored in navy blue while D797 belongs to Hel2 is colored in lemon) and T331I mutant (colored in dark blue).

Structural alignment of wild-type MDA5 and the T331I mutant monomers revealed that both constructs bind approximately 14 base pairs of dsRNA (Fig. S10A). This consistent RNA binding footprint suggests that the T331I mutation does not alter the overall architecture of the MDA5/dsRNA complex. However, the conformation of bound nucleotide differs between the two structures: in the T331I mutant, ATP is shifted outward from the nucleotide-binding pocket compared to AMPPNP in the wild-type structure (Fig. 5C). This observation aligns with our finding that fewer amino acids participate in stabilizing ATP in the T331I mutant (Fig. 4D). Additionally, we clearly observed a shift in the positions of two magnesium ions, further supporting the displacement of ATP. The associated water molecule also relocates to a different position. Direct comparison of the amino acids involved in nucleotide binding revealed a subtle conformational change in the P-loop, and the dissociation of residues D443, E444, Q371, and D797 from the nucleotide-binding pocket (Fig. 5D). Together, these results indicate that the T331I mutation alters both the flexibility of the filament and the configuration of the nucleotide-binding pocket, disrupting the precise positioning of ATP and thereby impairing ATP hydrolysis.

## DISCUSSION

Using an integrated set of orthogonal approaches that combine luciferase reporter assays, biochemical analyses and cryoEM structures, we establish that structural changes within the ATPase active site cause stable MDA5 complex assembly on short RNAs, resulting in subsequent dysregulation of the MDA5 receptor. Furthermore, our comparative analysis of MDA5 and the T331I mutant provides clearer mechanistic understanding of MDA5-RNA filament assembly and the impact of mutations on dysregulation of the MDA5 signaling pathway (Fig. S11). T331I likely increases the statistical likelihood of forming a signaling-competent complex on dsRNAs of any length, thereby increasing the probability that even very short dsRNAs can trigger signaling. The increased initiation rate, and possibly the enhanced persistence of T331I relative to WT may promote filament formation on both short and long dsRNAs.

Previous studies have demonstrated that MDA5 assembles cooperatively on dsRNA molecules, driving the formation of stable, well-ordered filaments that coat dsRNAs in a sequence-independent manner(41). In living cells, this powerful filament-forming assembly force is countered by an opposing force: ATP hydrolysis by MDA5 weakens its binding to dsRNA and promotes filament disassembly(24,37,38,42-44). This “push and pull” of cooperative protein loading and ATP-induced unloading or translocation enables wild-type MDA5 to behave with selectivity, forming stable RNA complexes capable of signaling only on very long dsRNAs such as viral replication intermediates while forming weak, dynamic complexes on shorter host dsRNAs in the cell(44) (Fig. S11, left). This selectivity for long dsRNAs is lost in MDA5 variants that display reduced ATPase activity or enhanced RNA binding affinity. Mutants such as R337G, M854K, T331I form stable complexes even on short host dsRNAs, leading to constitutive interferon signaling and inflammatory disease (Fig. S11, right)(13,26,44-46). Despite numerous excellent prior studies, significant gaps in our understanding of the MDA5 dysregulation process have remained. Previous work on disease-associated MDA5 mutants involved amino acids that are located far outside of the ATPase active site, leaving open the possibility that these induce specific pathological conformations of MDA5 that change protein stability or that they induce allosteric effects that cause MDA5 to lose selectivity for long dsRNA(26). Here we resolve this issue by explicitly disrupting the active-site interaction network between MDA5 T331I and ATP, which specifically blocks ATP hydrolysis and induces signaling on short dsRNAs (Fig. 1C and Fig. 1D). T331I might in fact be so sensitive to endogenous dsRNA that its expression alone might be sufficient to trigger signaling (Fig. S2). Consistent with previous studies on the GOF R337G variant, our findings establish a direct link between ATP hydrolysis and dsRNA length discrimination(44). A second issue is that stable, putatively functional MDA5 complexes on short dsRNAs have never been directly visualized structurally, thus their existence has remained hypothetical until now. Using cell-based reporter assays, we showed that the ATPase-deficient T331I mutant induces robust signaling on diverse dsRNA molecules that range from 30 to 500 base pairs in length (Fig. 1C). In parallel, electron microscopy studies on T331I enabled us to visualize active, MDA5-dsRNA signaling complexes on duplexes only 112bp in length (Fig. 3A and Fig. 4A). Taken together, these findings establish that, without the opposing effect of ATP hydrolysis, functional MDA5 complexes readily form on dsRNAs that are very short and within the range typical of abundant host RNAs. In addition, we establish that short filaments formed by wild-type MDA5 in the absence of ATP rapidly disassemble in the presence of ATP, while the T331I mutant, lacking ATPase activity, forms compact, stable filaments on short dsRNA molecules under all conditions (Fig. 2C and Fig. S4). Interestingly, recent work on the mechanism of LGP2 suggests that it serves a similar function: anchoring MDA5 filaments in a configuration that is sufficiently stable to form signaling-competent complexes even on short dsRNAs(44,47-49). These findings refine and extend the emerging view that the ATPase activity of MDA5 is important for proof-reading via dissociation or translocation of MDA5 on dsRNA. In wild□type MDA5, ATP hydrolysis destabilizes filaments on shorter duplexes by suppressing spontaneous filament assembly, thereby enforcing lengthfilament assembly, thereby enforcingdependent discrimination between viral and host RNAs. Consistent with a previously proposed translocation□based model(44), our structural and biochemical analyses of the T331I mutant provide a direct mechanistic view of MDA5 dysregulation. Direct disruption of the ATPase active site converts MDA5 from a dynamic protein into a static, filament□forming receptor that stably engages short dsRNA. This shift has fundamental implications for the physical characteristics of MDA5 signaling complexes, even in healthy cells.

For more than a decade, the observation that MDA5 interferon induction is exclusively stimulated by long dsRNA molecules (>500 bp) has been equated with the idea that the minimal MDA5 signaling complex is an extremely long MDA5-dsRNA filament, and that large prion-like complexes of MDA5 on RNA are required for productive interaction with MAVS(9,24,50). This generally-accepted model has always been inconsistent with the fact that long dsRNA duplexes are rare in human cells, and that MDA5 GOF mutants can be activated by short human dsRNAs such as rRNA fragments(51). By establishing that MDA5 can form stable, active signaling complexes on dsRNAs as short as 30 base-pairs, we demonstrate that long MDA5-RNA filaments are not a required form of the active MDA5 signaling complex, that kilobase-pair filaments are not specifically required for a functional interaction with MAVS(52) and signaling-competent MDA5-dsRNA complexes can be relatively small, as long as they are stable. The requirement for long double-stranded RNAs by wild-type MDA5 does not reflect structural requirements of the actual signaling complex on the mitochondrial membrane(24). Rather, it is due to the fact that only long RNA duplexes can support a central core of MDA5 oligomers capable of withstanding ATP-driven erosion(36), which degrades nascent filaments from both duplex ends, in a manner similar to that of ATP-induced disassembly at microtubule termini(37,53,54). Our findings, building upon previous work, demonstrate that MDA5 mutant-dsRNA signaling complexes can form compact oligomers containing as few as two MDA5 subunits, in addition to extended filaments. This suggests that MDA5 signaling complexes share more similarities with the small dsRNA-RIG-I signaling complexes than previously recognized, indicating the potential for additional commonalities in their mechanisms of MAVS activation and downstream events.

The cryoEM structures of the T331I MDA5/dsRNA complex with ATP and the wild-type MDA5/dsRNA complex with AMPPNP provide new insights into the chemical mechanism of RNA-dependent ATP hydrolysis in RLRs and other SF2 RNA helicases. Structural comparisons reveal that Thr331I forms a consistent helical twist (~72°) on short dsRNA, whereas wild-type MDA5 forms a more variable helical twist (~80°-82°), suggesting increased rigidity of the T331I mutant filament. The T331I mutant also exhibits a larger monomer-monomer interface compared to wild-type MDA5, suggesting enhanced filament stability, consistent with cellular luciferase assays showing that T331I can be robustly activated by short dsRNAs. Additionally, in the wild-type structure with AMPPNP, Thr331 forms polar contacts with the γ-phosphate, positioning it near Asp443 and Glu444 of the DECH motif, thereby facilitating nucleophilic attack by water. Our high-resolution cryoEM data thus offer a novel structural perspective on the role of T331I mutation in the helicase motif I, an evolutionary conserved region further underscoring the importance of modulating the ATP-dependent functions of MDA5. Although the overall architecture of the MDA5-dsRNA filament is similar in both structures, comparative analysis reveals a marked perturbation within the ATP binding pocket of the mutant that disrupts both ATP positioning and critical interactions necessary for hydrolysis. These findings elucidate how subtle alterations in the ATP binding site can directly affect SF2 enzyme function, highlighting the profound impact of a single amino acid substitution. Collectively, these insights advance our understanding of MDA5 mechanistic regulation, offering a foundation for the rational design of potent MDA5 agonists and antagonists as novel therapeutics for viral infections and autoimmune diseases.

## ACKNOWLEDGEMENTS

A.M.P. is an Investigator and L.X. was a Research Specialist of the Howard Hughes Medical Institute (HHMI) and now L.X. is an Assistant Professor at UF Scripps. We thank Shenping Wu, Marc Llaguno, Jianfeng Lin, and Kaifeng Zhou at Yale CryoEM Facility for assistance in cryoEM data collection. We thank Tianshuo Liu, Junhui Peng, Swapnil Devarkar, Shuhui Wang and Long Han for insightful discussion about cryoEM data processing and model building. This work was supported by HHMI and Yale Colton Center. CryoEM data were collected with microscopes at the Yale CryoEM Resource Core that is funded in part by the NIH (S10OD023603).

## AUTHOR CONTRIBUTIONS

L.X. and A.M.P. conceived the project and designed the experiments. L.X. and A. P. performed biochemical purification of the protein. L. X. prepared cryoEM samples. J. H. performed RT-qPCR and conducted data analysis. L.X., and K.C. performed cryoEM data analysis and model building. L.X., K.C. and A.M.P. contributed to the structure analysis. R.G. and A.P. assisted in biochemical characterization of ATPase activity. L.X. and A.M.P wrote the manuscript.

## SUPPLEMENTARY DATA

Supplementary Figs 1 to 11

Supplementary Tables 1 to 2

Supplementary Data are available at NAR online.

## DATA AVAILABILITY

All data needed to evaluate the conclusions in the paper are present in the paper and/or the Supplementary Materials. CryoEM maps generated in this study are deposited in the Electron Microscopy Data Bank with codes EMD-72484 (MonoMDA5/dsRNA), EMD-72485 (diMDA5/dsRNA), EMD-72284 (TetraMDA5/dsRNA), EMD-72244 (MonoT331I/dsRNA),

EMD-72243 (diT331I/dsRNA), EMD-72285 (TetraT331I/dsRNA). Model coordinates have been deposited in the Protein Data Bank (PDB) under DOIs: https://doi.org/10.2210/pdb9Y4J/pdb (MonoMDA5/dsRNA), https://doi.org/10.2210/pdb9Y4L/pdb (diMDA5/dsRNA), https://doi.org/10.2210/pdb9Q60/pdb (monoT331I/dsRNA), https://doi.org/10.2210/pdb9Q5W/pdb (diT331I/dsRNA).

## REFERENCES

1. Triantafilou, K., Vakakis, E., Kar, S., Richer, E., Evans, G.L. and Triantafilou, M. (2012) Visualisation of direct interaction of MDA5 and the dsRNA replicative intermediate form of positive strand RNA viruses. J Cell Sci, 125, 4761–4769.

2. Dias Junior, A.G., Sampaio, N.G. and Rehwinkel, J. (2019) A Balancing Act: MDA5 in Antiviral Immunity and Autoinflammation. Trends Microbiol, 27, 75–85.

3. Cao, X., Ding, Q., Lu, J., Tao, W., Huang, B., Zhao, Y., Niu, J., Liu, Y.J. and Zhong, J. (2015) MDA5 plays a critical role in interferon response during hepatitis C virus infection. J Hepatol, 62, 771–778.

4. Pulido, M.R., Martinez-Salas, E., Sobrino, F. and Saiz, M. (2020) MDA5 cleavage by the Leader protease of foot-and-mouth disease virus reveals its pleiotropic effect against the host antiviral response. Cell Death Dis, 11, 718.

5. Rebendenne, A., Valadao, A.L.C., Tauziet, M., Maarifi, G., Bonaventure, B., McKellar, J., Planes, R., Nisole, S., Arnaud-Arnould, M., Moncorge, O. et al. (2021) SARS-CoV-2 triggers an MDA-5-dependent interferon response which is unable to control replication in lung epithelial cells. J Virol, 95.

6. Yin, X., Riva, L., Pu, Y., Martin-Sancho, L., Kanamune, J., Yamamoto, Y., Sakai, K., Gotoh, S., Miorin, L., De Jesus, P.D. et al. (2021) MDA5 Governs the Innate Immune Response to SARS-CoV-2 in Lung Epithelial Cells. ell Rep, 34, 108628.

7. Sampaio, N.G., Chauveau, L., Hertzog, J., Bridgeman, A., Fowler, G., Moonen, J.P., Dupont, M., Russell, R.A., Noerenberg, M. and Rehwinkel, J. (2021) The RNA sensor MDA5 detects SARS-CoV-2 infection. Sci Rep, 11, 13638.

8. Thorne, L.G., Reuschl, A.K., Zuliani-Alvarez, L., Whelan, M.V.X., Turner, J., Noursadeghi, M., Jolly, C. and Towers, G.J. (2021) SARS-CoV-2 sensing by RIG-I and MDA5 links epithelial infection to macrophage inflammation. EMBO J, 40, e107826.

9. Peisley, A., Jo, M.H., Lin, C., Wu, B., Orme-Johnson, M., Walz, T., Hohng, S. and Hur, S. (2012) Kinetic mechanism for viral dsRNA length discrimination by MDA5 filaments. Proc Natl Acad Sci U S A, 109, E3340–3349.

10. Rawling, D.C., Fitzgerald, M.E. and Pyle, A.M. (2015) Establishing the role of ATP for the function of the RIG-I innate immune sensor. Elife, 4.

11. Singh, R., Wu, Y., del Valle, A.H., Leigh, K.E., Mong, S., Cheng, M.T.K., Ferguson, B.J. and Modis, Y. (2024) Contrasting functions of ATP hydrolysis by MDA5 and LGP2 in viral RNA sensing. J Biol Chem, 300.

12. Quack, S., Maity, S., America, P.P.B., Klein, M., Herrero Del Valle, A., Singh, R., Joiner, J.D., Rashid, Z.M., Smitskamp, Q., Rivas, P. et al. (2026) MDA5 generates compact ribonucleoprotein complexes via ATP-dependent double-stranded RNA unwinding. Nucleic Acids Res, 54.

13. Rice, G.I., Del Toro Duany, Y., Jenkinson, E.M., Forte, G.M., Anderson, B.H., Ariaudo, G., Bader-Meunier, B., Baildam, E.M., Battini, R., Beresford, M.W. et al. (2014) Gain-of-function mutations in IFIH1 cause a spectrum of human disease phenotypes associated with upregulated type I interferon signaling. Nat Genet, 46, 503–509.

14. Ohto, T., Tayeh, A.A., Nishikomori, R., Abe, H., Hashimoto, K., Baba, S., Arias-Loza, A.P., Soda, N., Satoh, S., Matsuda, M. et al. (2022) Intracellular virus sensor MDA5 mutation develops autoimmune myocarditis and nephritis. J Autoimmun, 127, 102794.

15. Funabiki, M., Kato, H., Miyachi, Y., Toki, H., Motegi, H., Inoue, M., Minowa, O., Yoshida, A., Deguchi, K., Sato, H. et al. (2014) Autoimmune disorders associated with gain of function of the intracellular sensor MDA5. Immunity, 40, 199–212.

16. Miner, J.J. and Diamond, M.S. (2014) MDA5 and autoimmune disease. Nat Genet, 46, 418–419.

17. Sadler, A.J. (2018) The role of MDA5 in the development of autoimmune disease. J Leukoc Biol, 103, 185–192.

18. Najm, R., Yavuz, L., Jain, R., El Naofal, M., Ramaswamy, S., Abuhammour, W., Loney, T., Nowotny, N., Alsheikh-Ali, A., Abou Tayoun, A. et al. (2024) IFIH1 loss of function predisposes to inflammatory and SARS-CoV-2-related infectious diseases. Scand J Immunol, 100, e13373.

19. Rice, G.I., Park, S., Gavazzi, F., Adang, L.A., Ayuk, L.A., Van Eyck, L., Seabra, L., Barrea, C., Battini, R., Belot, A. et al. (2020) Genetic and phenotypic spectrum associated with IFIH1 gain-of-function. Hum Mutat, 41, 837–849.

20. Rutsch, F., MacDougall, M., Lu, C., Buers, I., Mamaeva, O., Nitschke, Y., Rice, G.I., Erlandsen, H., Kehl, H.G., Thiele, H. et al. (2015) A specific IFIH1 gain-of-function mutation causes Singleton-Merten syndrome. Am J Hum Genet, 96, 275–282.

21. Bursztejn, A.C., Briggs, T.A., del Toro Duany, Y., Anderson, B.H., O’Sullivan, J., Williams, S.G., Bodemer, C., Fraitag, S., Gebhard, F., Leheup, B. et al. (2015) Unusual cutaneous features associated with a heterozygous gain-of-function mutation in IFIH1: overlap between Aicardi-Goutieres and Singleton-Merten syndromes. Br J Dermatol, 173, 1505–1513.

22. de Carvalho, L.M., Ngoumou, G., Park, J.W., Ehmke, N., Deigendesch, N., Kitabayashi, N., Melki, I., Souza, F.F.L., Tzschach, A., Nogueira-Barbosa, M.H. et al. (2017) Musculoskeletal Disease in MDA5-Related Type I Interferonopathy: A Mendelian Mimic of Jaccoud’s Arthropathy. Arthritis Rheumatol, 69, 2081–2091.

23. Pyle, A.M. (2008) Translocation and unwinding mechanisms of RNA and DNA helicases. Annu Rev Biophys, 37, 317–336.

24. Peisley, A., Lin, C., Wu, B., Orme-Johnson, M., Liu, M., Walz, T. and Hur, S. (2011) Cooperative assembly and dynamic disassembly of MDA5 filaments for viral dsRNA recognition. Proc Natl Acad Sci U S A, 108, 21010–21015.

25. Singh, R., Wu, Y., Herrero Del Valle, A., Leigh, K.E., Mong, S., Cheng, M.T.K., Ferguson, B.J. and Modis, Y. (2024) Contrasting functions of ATP hydrolysis by MDA5 and LGP2 in viral RNA sensing. J Biol Chem, 300, 105711.

26. Yu, Q., Herrero Del Valle, A., Singh, R. and Modis, Y. (2021) MDA5 disease variant M854K prevents ATP-dependent structural discrimination of viral and cellular RNA. Nat Commun, 12, 6668.

27. Guo, R. and Pyle, A.M. (2022) Monitoring functional RNA binding of RNA-dependent ATPase enzymes such as SF2 helicases using RNA dependent ATPase assays: A RIG-I case study. Helicase Enzymes, Pt B, 673, 39–52.

28. Varadi, M., Anyango, S., Deshpande, M., Nair, S., Natassia, C., Yordanova, G., Yuan, D., Stroe, O., Wood, G., Laydon, A. et al. (2022) AlphaFold Protein Structure Database: massively expanding the structural coverage of protein-sequence space with high-accuracy models. Nucleic Acids Res, 50, D439–D444.

29. Jumper, J., Evans, R., Pritzel, A., Green, T., Figurnov, M., Ronneberger, O., Tunyasuvunakool, K., Bates, R., Zidek, A., Potapenko, A. et al. (2021) Highly accurate protein structure prediction with AlphaFold. Nature, 596, 583–589.

30. Goddard, T.D., Huang, C.C. and Ferrin, T.E. (2007) Visualizing density maps with UCSF Chimera. J Struct Biol, 157, 281–287.

31. Afonine, P.V., Poon, B.K., Read, R.J., Sobolev, O.V., Terwilliger, T.C., Urzhumtsev, A. and Adams, P.D. (2018) Real-space refinement in PHENIX for cryo-EM and crystallography. Acta Crystallogr D Struct Biol, 74, 531–544.

32. Adams, P.D., Afonine, P.V., Bunkoczi, G., Chen, V.B., Davis, I.W., Echols, N., Headd, J.J., Hung, L.W., Kapral, G.J., Grosse-Kunstleve, R.W. et al. (2010) PHENIX: a comprehensive Python-based system for macromolecular structure solution. Acta Crystallogr D Biol Crystallogr, 66, 213–221.

33. Pettersen, E.F., Goddard, T.D., Huang, C.C., Meng, E.C., Couch, G.S., Croll, T.I., Morris, J.H. and Ferrin, T.E. (2021) UCSF ChimeraX: Structure visualization for researchers, educators, and developers. Protein Sci, 30, 70–82.

34. Linehan, M.M., Dickey, T.H., Molinari, E.S., Fitzgerald, M.E., Potapova, O., Iwasaki, A. and Pyle, A.M. (2018) A minimal RNA ligand for potent RIG-I activation in living mice. Sci Adv, 4, e1701854.

35. Thoresen, D.T., Galls, D., Gotte, B., Wang, W. and Pyle, A.M. (2023) A rapid RIG-I signaling relay mediates efficient antiviral response. Mol Cell, 83, 90–104 e104.

36. Berke, I.C. and Modis, Y. (2012) MDA5 cooperatively forms dimers and ATP-sensitive filaments upon binding double-stranded RNA. EMBO J, 31, 1714–1726.

37. Wu, B., Peisley, A., Richards, C., Yao, H., Zeng, X., Lin, C., Chu, F., Walz, T. and Hur, S. (2013) Structural basis for dsRNA recognition, filament formation, and antiviral signal activation by MDA5. Cell, 152, 276–289.

38. Yu, Q., Qu, K. and Modis, Y. (2018) Cryo-EM Structures of MDA5-dsRNA Filaments at Different Stages of ATP Hydrolysis. Mol Cell, 72, 999–1012 e1016.

39. Krissinel, E. and Henrick, K. (2007) Inference of macromolecular assemblies from crystalline state. J Mol Biol, 372, 774–797.

40. Tauchert, M.J., Fourmann, J.B., Lührmann, R. and Ficner, R. (2017) Structural insights into the mechanism of the DEAH-box RNA helicase Prp43. Elife, 6.

41. Fitzgerald, M.E., Vela, A. and Pyle, A.M. (2014) Dicer-related helicase 3 forms an obligate dimer for recognizing 22G-RNA. Nucleic Acids Res, 42, 3919–3930.

42. Kato, H., Takeuchi, O., Mikamo-Satoh, E., Hirai, R., Kawai, T., Matsushita, K., Hiiragi, A., Dermody, T.S., Fujita, T. and Akira, S. (2008) Length-dependent recognition of double-stranded ribonucleic acids by retinoic acid-inducible gene-I and melanoma differentiation-associated gene 5. J Exp Med, 205, 1601–1610.

43. Peisley, A., Jo, M.H., Lin, C., Wu, B., Orme-Johnson, M., Walz, T., Hohng, S. and Hur, S. (2012) Kinetic mechanism for viral dsRNA length discrimination by MDA5 filaments. P Natl Acad Sci USA, 109, E3340–E3349.

44. Han, X.P., Rao, M., Chang, Y., Zhu, J.Y., Cheng, J., Li, Y.T., Qiong, W., Ye, S.C., Zhang, Q., Zhang, S.Q. et al. (2025) ATP-dependent one-dimensional movement maintains immune homeostasis by suppressing spontaneous MDA5 filament assembly. Cell Res, 35, 900–912.

45. Wallace, C., Singh, R. and Modis, Y. (2025) MDA5 variants trade antiviral activity for protection from autoimmune disease. BMC Med Genomics, 18, 101.

46. Singh, R., Joiner, J.D., Herrero Del Valle, A., Zwaagstra, M., Jobe, I., Ferguson, B.J., van Kuppeveld, F.J.M. and Modis, Y. (2025) Molecular basis of autoimmune disease protection by MDA5 variants. Cell Rep, 44, 115754.

47. Kurihara, N., Isayama, Y., Zhang, J., Yamashita, T., Awaji, K., Ito, Y., Yoshizaki, A., Kouwaki, T., Oshiumi, H., Nishimasu, H. et al. (2026) Molecular mechanism of MDA5 nucleation and filament formation by LGP2. Mol Cell, 86, 707–721 e705.

48. Jang, J. and Jo, M.H. (2025) LGP2 stops MDA5 translocation to start antiviral signaling. Cell Res, 35, 785–786.

49. Uchikawa, E., Lethier, M., Malet, H., Brunel, J., Gerlier, D. and Cusack, S. (2016) Structural Analysis of dsRNA Binding to Anti-viral Pattern Recognition Receptors LGP2 and MDA5. Mol Cell, 62, 586–602.

50. Duany, Y.D., Wu, B. and Hur, S. (2015) MDA5-filament, dynamics and disease. Curr Opin Virol, 12, 20–25.

51. Ahmad, S., Mu, X., Yang, F., Greenwald, E., Park, J.W., Jacob, E., Zhang, C.Z. and Hur, S. (2018) Breaching Self-Tolerance to Alu Duplex RNA Underlies MDA5-Mediated Inflammation. Cell, 172, 797–810 e713.

52. Vazquez, C. and Horner, S.M. (2015) MAVS Coordination of Antiviral Innate Immunity. J Virol, 89, 6974–6977.

53. Bershadsky, A.D. and Gelfand, V.I. (1981) ATP-dependent regulation of cytoplasmic microtubule disassembly. Proc Natl Acad Sci U S A, 78, 3610–3613.

54. Hartman, J.J. and Vale, R.D. (1999) Microtubule disassembly by ATP-dependent oligomerization of the AAA enzyme katanin. Science, 286, 782–785.

