## Supplementary Figures and Tables for "Establishing the molecular basis for MDA5 mutation-linked autoimmunity"

**The PDF file includes:**

**Fig. S1 to S11**

**Table S1 to S2**

**Figure S1**

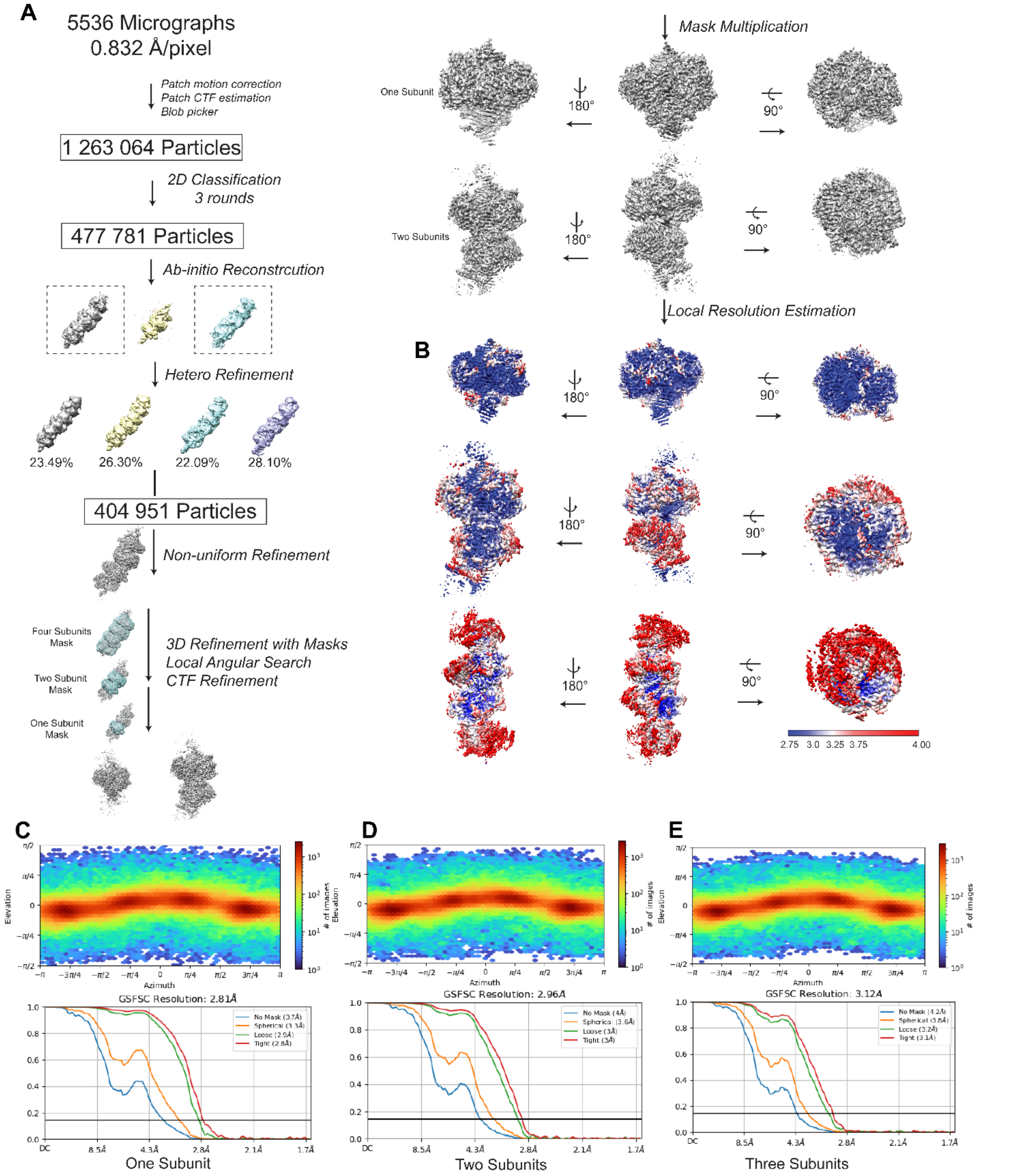


**Figure S1. CryoEM workflow for the wild-type MDA5/dsRNA in presence of AMPPNP.** A. CryoEM data processing workflow for the wild-type MDA5/dsRNA in presence of AMPPNP. B. Local resolution map of the wild-type MDA5/dsRNA in presence of AMPPNP with one subunit (top panel) and two subunits on the dsRNA (bottom panel). C-E. Particle orientation distribution of one, two and four protein subunit on dsRNA (top panel). FSC curve for the cryoEM reconstruction of one, two and four protein subunit on dsRNA (bottom panel).

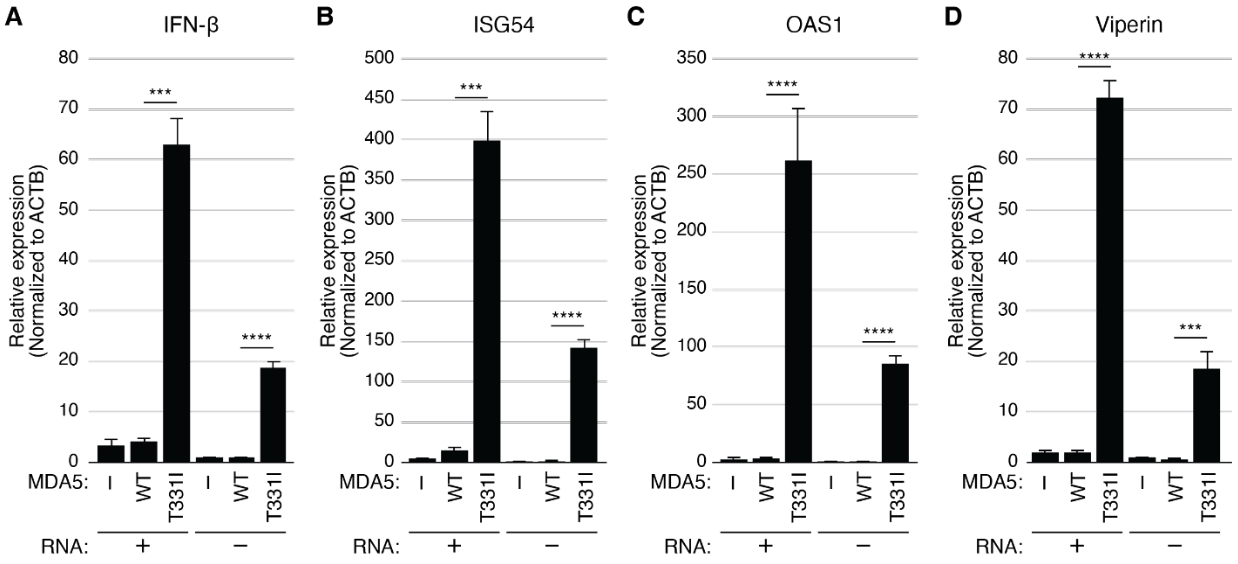


**Figure S2. RT-qPCR of IFN response genes in cells expressing wild-type MDA5 and T331I mutant.** A-D. RT-qPCR analysis of IFN-β (A), ISG54 (B), OAS1 (C), and Viperin (D). Cells were prepared under the same transfection conditions as those used for the luciferase assay. Cells were stimulated by poly (I:C). Bar graphs represent the average of three biological replicates, and error bars indicate the standard error of the mean. P values were calculated using an unpaired two-tailed Student's t-test. *P < 0.05, **P < 0.01, ***P < 0.001, and ****P < 0.0001.

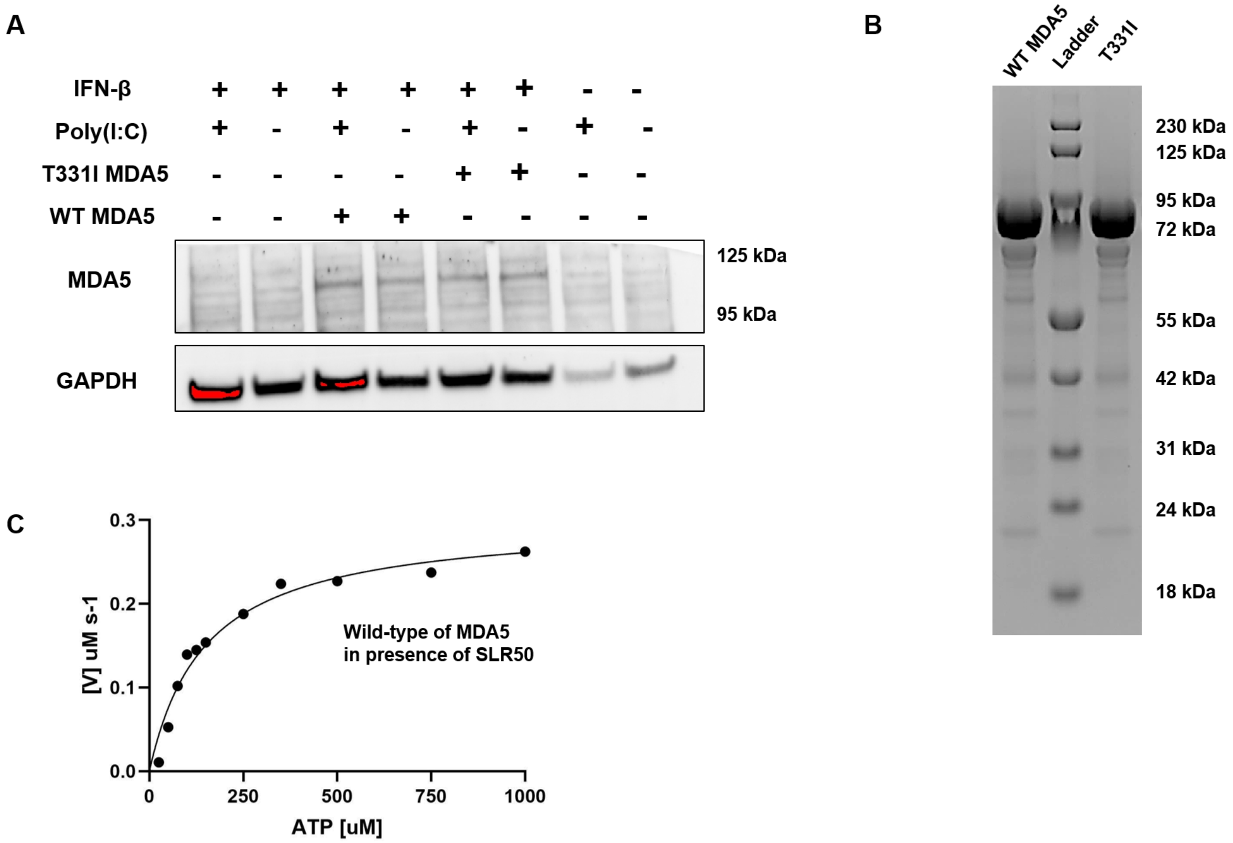


**Figure S3. Biochemistry data of MDA5 and T331I mutant.** A. Expression levels of MDA5, T331I and GAPDH were analyzed by immunoblotting with specific antibodies. Western blot of wild-type MDA5 and T331I mutant used samples from the luciferase assay. Mouse anti-MDA5 antibody was used to detect the expression of MDA5. **B.** SDS-PAGE for purified wild-type MDA5 and T331I mutant protein. 2uM protein was loaded. The gel was stained by Coomassie staining buffer. C. ATPase assay of wild-type MDA5 in presence SLR50.

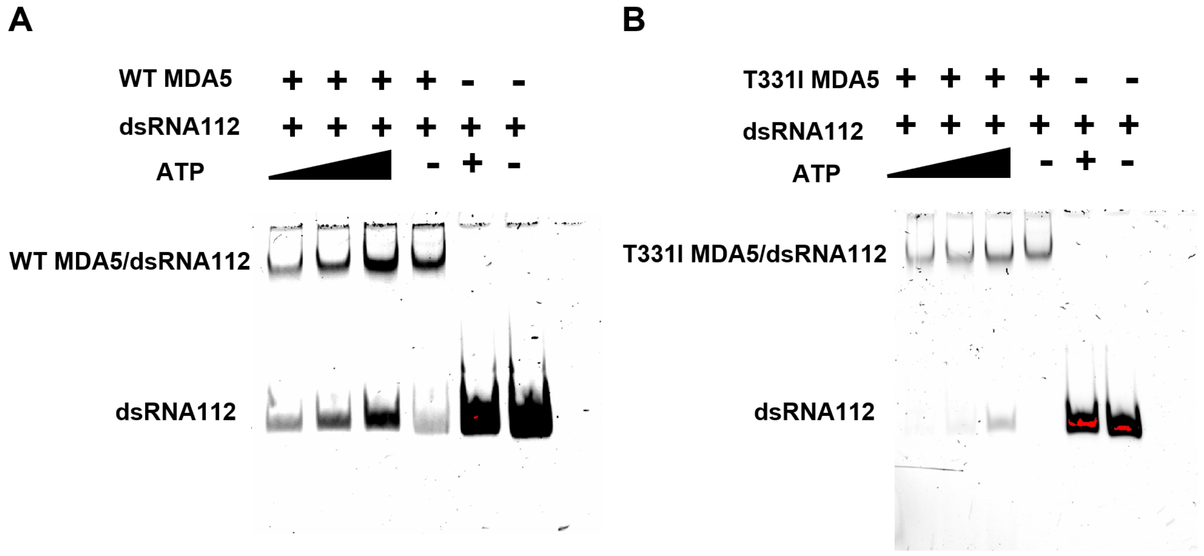


**Figure S4. EMSA for wild-type MDA5 and T331I with dsRNA in presence of ATP.** In the EMSA experiment, 20 nM RNA was incubated with 0.5 μM protein at room temperature for 30 minutes to allow complex formation. ATP was then added to final concentrations of 1 mM, 3 mM, or 5 mM, followed by an additional 6-minute incubation at room temperature. Subsequently, 2 μL of loading buffer was mixed with each 10 μL sample, and the mixtures were loaded onto an SDS-PAGE gel and stained with GelRed.

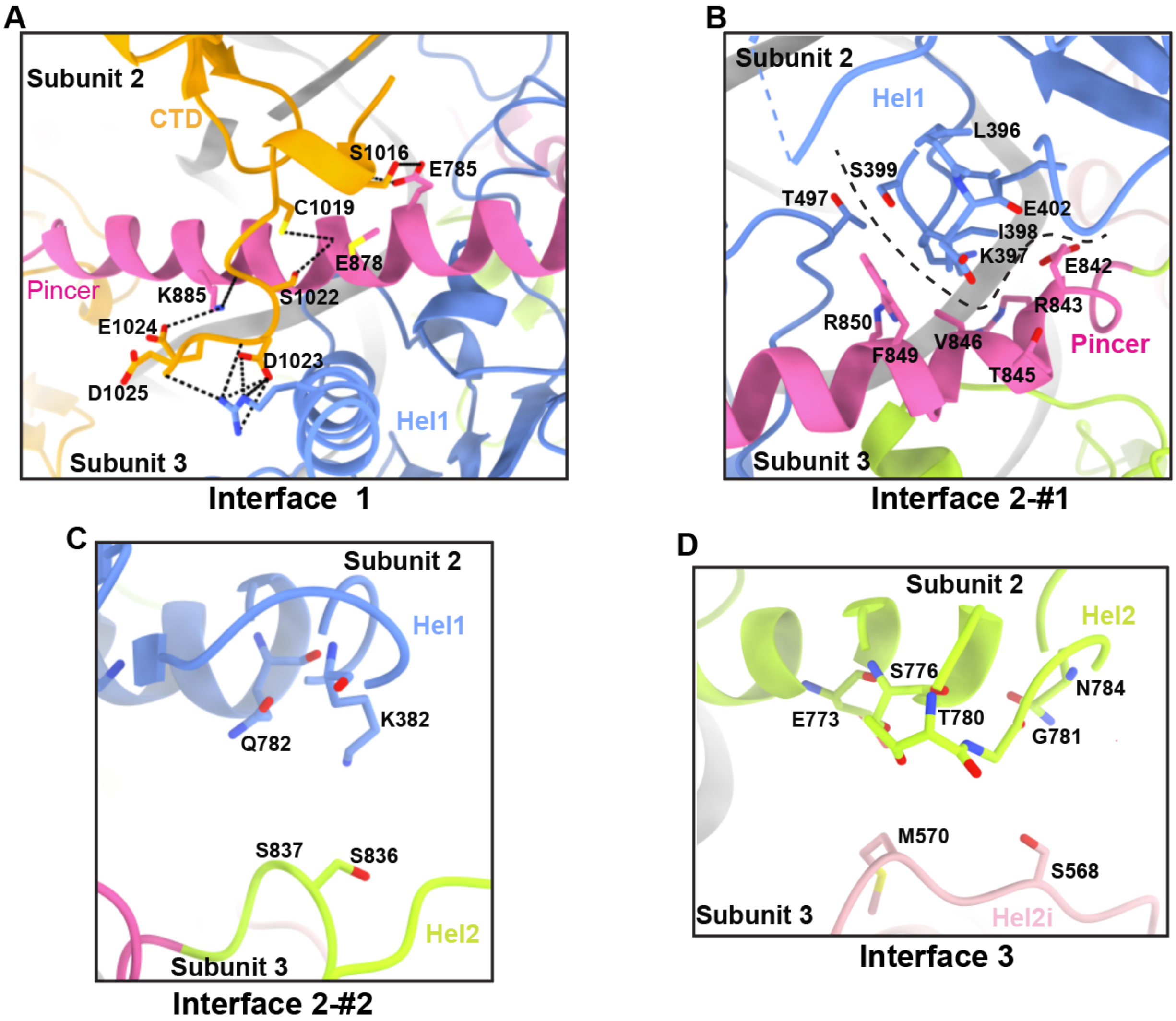


**Figure S5. Monomer-monomer interface of the wild-type MDA5/dsRNA.** A. CTD tail from subunit 2 (orange) forms extensive hydrophobic interactions and several hydrogen bonds (indicated with dash lines) with amino acids from the Pincer (pink) and Hel1of subunit 3 (blue). B-C. Amino acids from Hel1 of subunit 2 form extensive hydrophobic interactions with amino acids of Pincer and Hel2 of subunit 3. The dashed line indicates a groove that is formed by the Pincer and Hel2 of subunit3. D. Amino acids from Hel2 of subunit 2 forms hydrophobic interactions with amino acids of Hel2i of subunit 3.

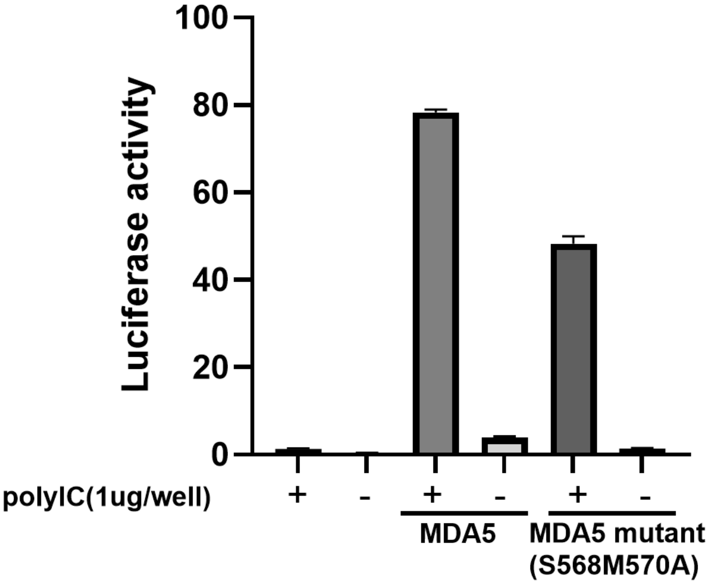


**Figure S6. IFN-β induction by MDA5 and the MDA5 monomer-monomer interface mutant MDA5(S568M570A)**. Upon poly(I:C) stimulation in HEK293T cells. Cells were transfected with 1µg/well poly(I:C). Luciferase activities were measured 12 h post-transfection. Data represent the mean ± SD of three technical replicates from one representative biological replicate. Statistical significance: **** $p < 0.0001$; by ordinary one-way ANOVA for multiple comparisons.

**Figure S7**

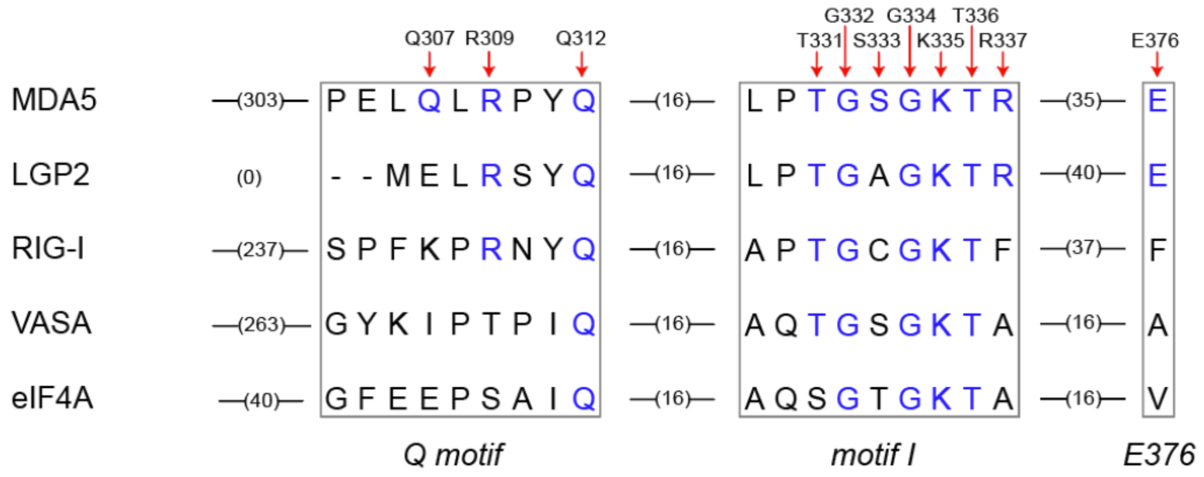


**Figure S7.** Multiple sequence alignment of RLR family members (human MDA5, LGP2, RIG-I) and related RNA helicases (human eIF4A and Drosophila VASA). Conserved amino acids are highlighted in blue.

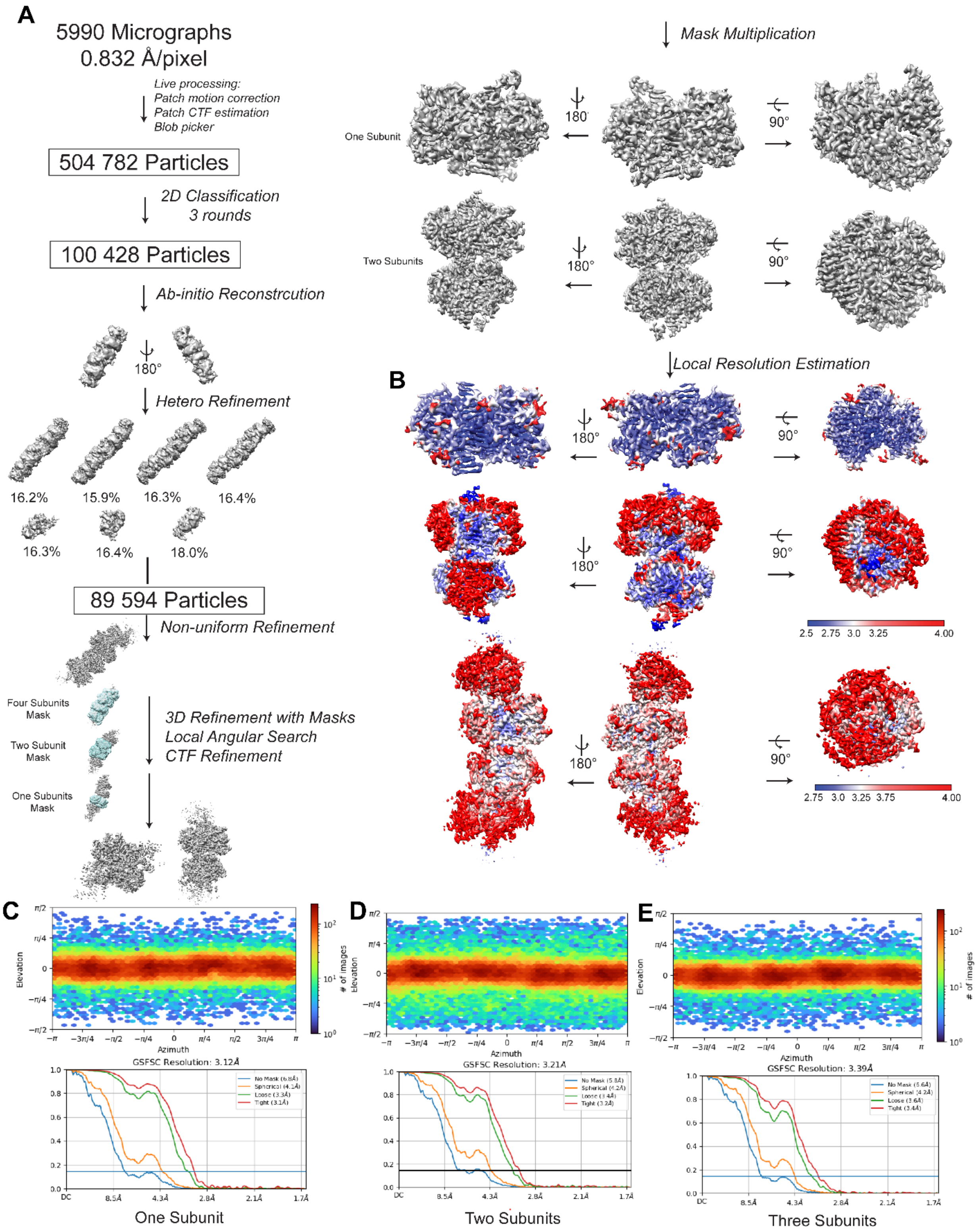


**Figure S8. CryoEM workflow for T331I/dsRNA in presence of ATP.** A. CryoEM data processing workflow for T331I/dsRNA in presence of ATP. B. Local resolution map of the T331I/dsRNA in presence of ATP with one, two and four subunits on the dsRNA. C-E. Particle orientation distribution and FSC curve for the cryoEM reconstruction of one, two and four protein subunits on dsRNA.

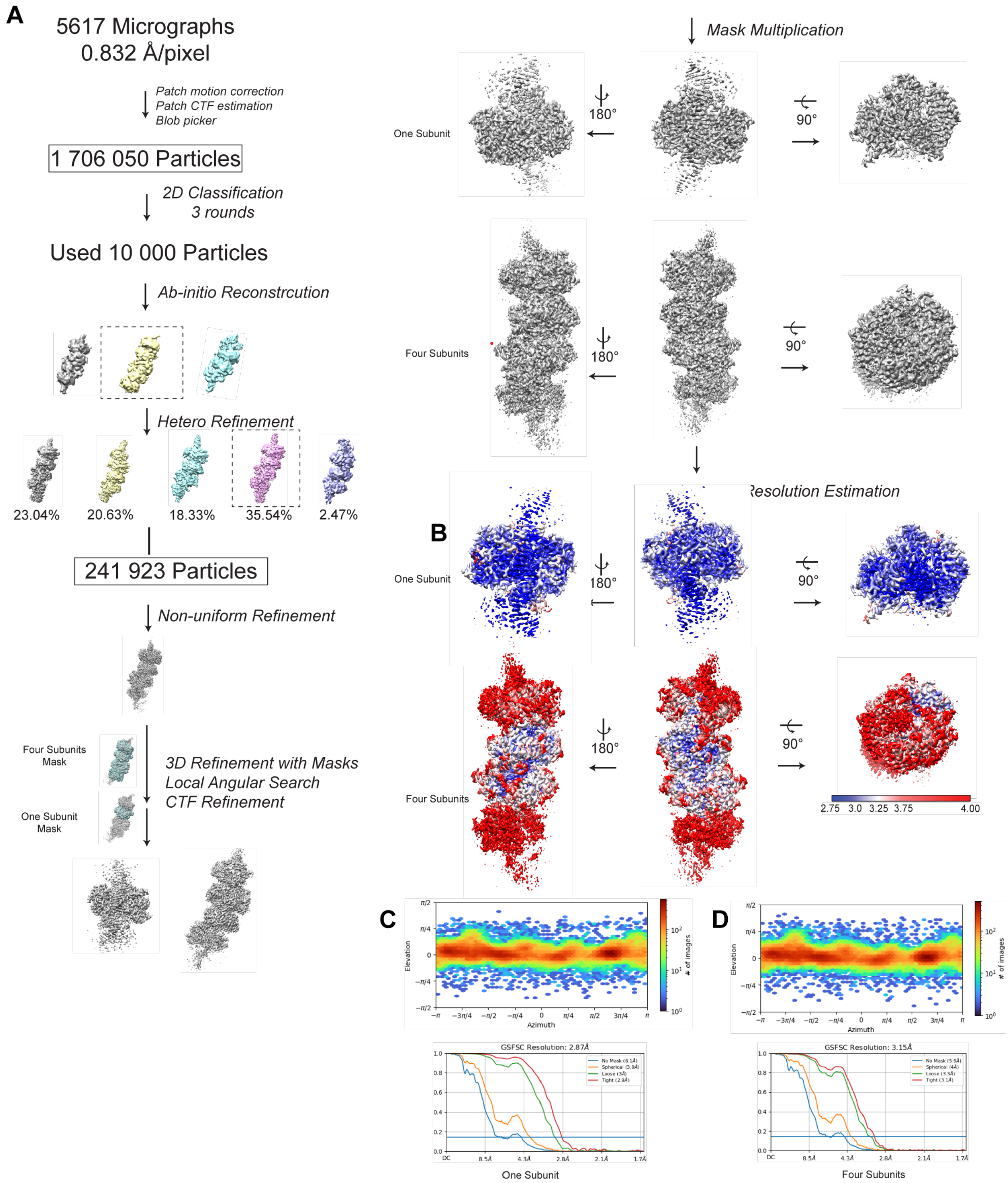


**Figure S9. CryoEM workflow for T331I/dsRNA in presence of AMPPNP.** A. CryoEM data processing workflow for T331I/dsRNA in presence of AMPPNP. B. Local resolution map of the T331I/dsRNA in presence of AMPPNP with one and four subunits on the dsRNA. C-D. Particle orientation distribution and FSC curve for the cryoEM reconstruction of one and four protein subunits on dsRNA.

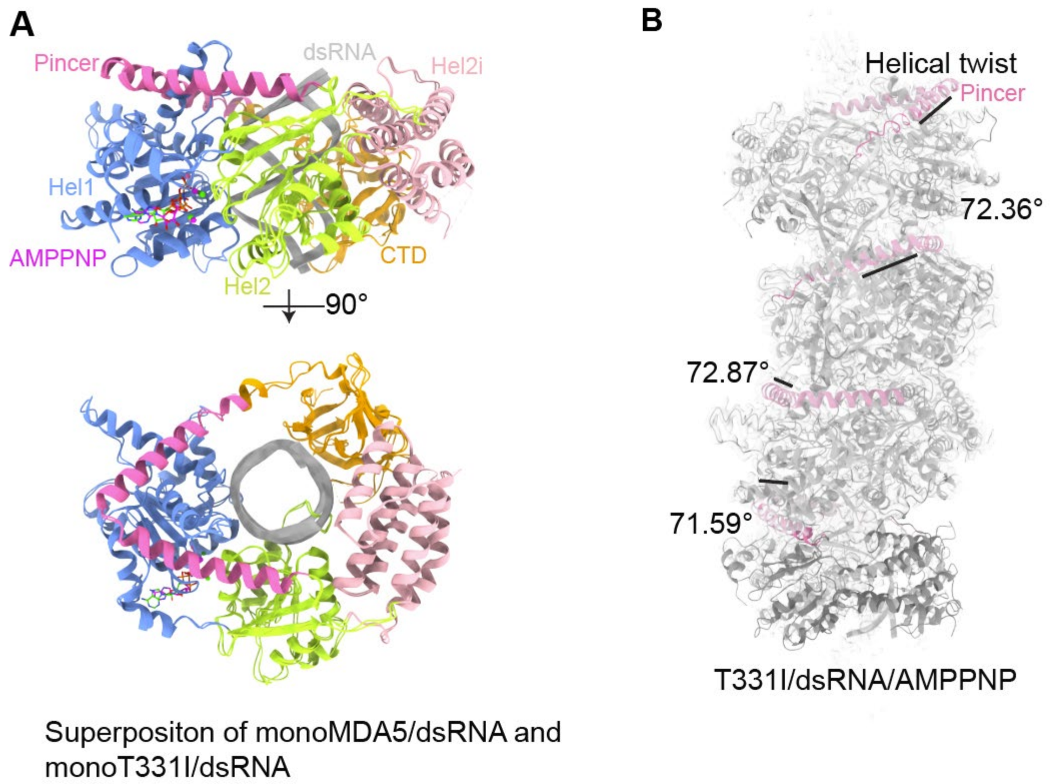


**Figure S10. Structure comparison of wild-type MDA5/dsRNA and T331I mutant/dsRNA.** A. Superposition of monoMDA5/dsRNA and monoT331I/dsRNA. The side view and top view of the superposition of monoMDA5/dsRNA and monoT331I/dsRNA structure, with ATP and AMPPNP shown as stick representations. B. The T331I mutant forms filaments with a constant helical twist on short dsRNA in presence of AMPPNP. The measured helical twist angles between subunits are 72.36° (subunit 1 to 2), 72.87° (subunit 2 to 3), and 71.59° (subunit 3 to 4).

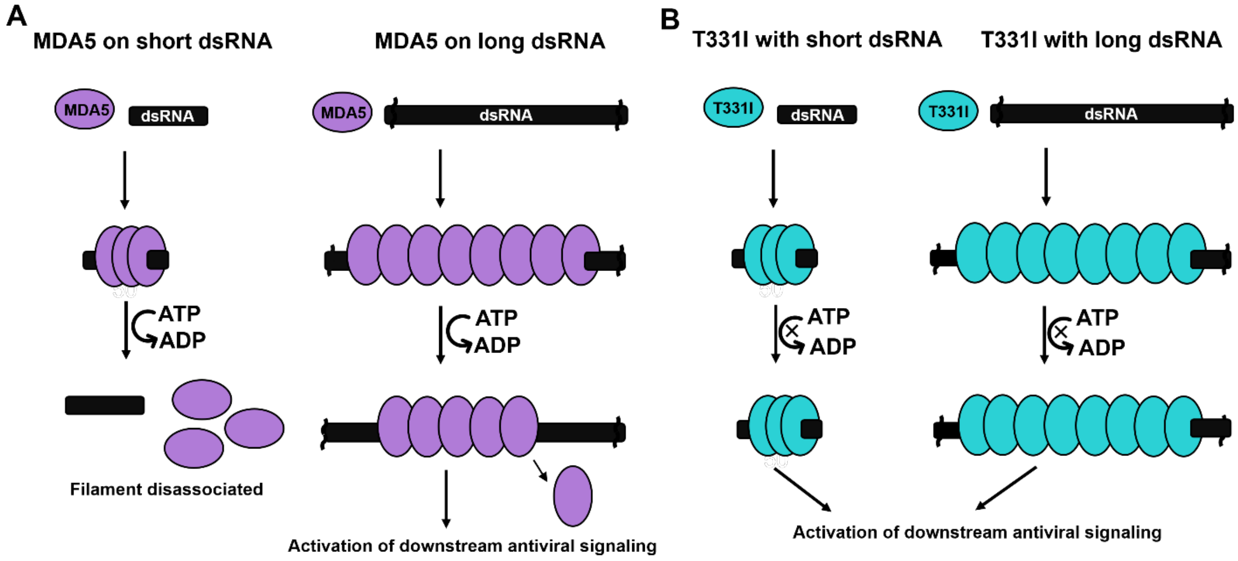


**Figure S11. Proposed model for MDA5 behavior on short and long dsRNA.** A. Wild-type MDA5 (purple). On short dsRNA (black bar): filament readily disassemble upon ATP hydrolysis. On long dsRNA: Filaments are more stable. Despite partially disassembled through ATP hydrolysis, sufficient MDA5 remains to maintain downstream antiviral signaling. B. MDA5 T331I mutant (cyan), on both short and long dsRNA: loss of ATPase activity prevents filament disassembly following ATP hydrolysis, resulting in stable filaments and sustained antiviral signaling.

There are several reasons that a WT MDA filament is much more likely to reach a signal-competent length on a long dsRNA than on a short dsRNA under conditions of abundant ATP. But two statistical reasons are straightforward to understand intuitively. As a starting premise, it is helpful to think about MDA5 signaling as a game, in which individual MDA5-RNA filaments are competing with one another, attempting to resist ATP-dependent erosion and survive long enough to engage with MAVS and induce a signal (which counts as a "win"). Imagine that there is a minimal filament size that is needed for engagement with MAVS (let's say 4 MDA5 units on a given dsRNA). Indeed, it is true that "winning" filaments of this size can form on both short and long dsRNAs, but those that form on long dsRNAs have two major advantages in the case of WT MDA5.

1**. Initiation**: Longer dsRNAs contain more nucleation sites for MDA5 binding and filament initiation than a comparable short RNA at the same molar concentration. Longer dsRNAs therefore have a higher statistical likelihood of forming a "winning" filament at some point along their length. For example, an RNA of 60 base pairs only has one opportunity, but an RNA of 300 base pairs has five (assuming an MDA5 site size of 15 bp and a signaling-competent minimal length of 4MDA5's on dsRNA).

2. **Persistence**: MDA5 units in the center of a filament form three extensive molecular interfaces that stabilize their position on the RNA: an interface with RNA and an interface with each of the two neighboring MDA5 molecules. By contrast, MDA5 units at a dsRNA terminus have only two interfaces (one with RNA and one with the flanking MDA5 molecule), and are therefore more likely to dissociate. The longer a filament is, the more MDA5 units it internalizes within the center and the more capable it will be of sustaining a "winning" length of at least 4MDA5 molecules, even as the ends erode. Long filaments can only form on long dsRNAs. There are no protective advantages to filaments that form on short dsRNAs.

**Table S1. Cryo-EM data collection, refinement, and validation statistics.**

**Table S1. Cryo-EM data collection, refinement, and validation statistics.**

| **Sample** | **Wild-type MDA5/dsRNA/AMPPNP** | | | **T331I/dsRNA/ATP** | | | **T331I/dsRNA/AMPPNP** | |
|---|---|---|---|---|---|---|---|---|
| **Data collection and processing** | | | | | | | | |
| Voltage (kV) | 300 | | | 300 | | | 300 | |
| Camera | K3 | | | K3 | | | K3 | |
| Symmetry imposed | C1 | | | C1 | | | C1 | |
| Magnification | 105,000x | | | 105,000x | | | 105,000x | |
| Defocus range (µm) | -0.5 to -2.0 | | | -0.5 to -2.0 | | | -0.5 to -2.0 | |
| Total dose (e-/$A^2$) | 51.98 | | | 50.49 | | | 59.02 | |
| Pixel size (Å) | 0.832 | | | 0.832 | | | 0.832 | |
| Micrographs collected | 5640 | | | 5990 | | | 6325 | |
| Micrographs processed | 3704 | | | 5989 | | | 5617 | |
| Initial particles | 646803 | | | 504782 | | | 1706050 | |
| **Refinement** | | | | | | | | |
| | MonoMDA5/dsRNA (one protein subunit on dsRNA) | DiMDA5/dsRNA (two protein subunits on dsRNA) | TetraMDA5/dsRNA (four protein subunits on dsRNA) | MonoT331I/dsRNA (one protein subunit on dsRNA) | DiT331I/dsRNA (two protein subunits on dsRNA) | TetraT331I/dsRNA (four protein subunits on dsRNA) | one protein subunit on dsRNA in presence of AMPPNP | four protein subunits on dsRNA in presence of AMPPNP |
| Particles | 402,040 | 402,040 | 404,951 | 75,762 | 89,594 | 79,446 | 83,677 | 85,865 |
| Map resolution (Å) | 2.81 | 2.96 | 3.04 | 3.12 | 3.21 | 3.39 | 2.87 | 3.15 |
| FSC threshold | 0.143 | 0.143 | 0.143 | 0.143 | 0.143 | 0.143 | 0.143 | 0.143 |
| Map sharpening B-factor ($A^2$) | -76.7 | -70.5 | -86.4 | -65 | -63.7 | -59.1 | -63 | -49 |
| **Model composition** | | | | | | | | |
| Non-hydrogen atoms | 5,943 | 12,108 | NA[1] | 6,014 | 12,242 | NA[1] | NA[3] | NA[1] |
| Protein residues | 664 | 1,336 | NA[1] | 673 | 1358 | NA[1] | NA[3] | NA[1] |
| RNA nucleotides | 26 | 60 | NA[1] | 26 | 60 | NA[1] | NA[3] | NA[1] |
| Ligands | AMPPNP | AMPPNP | NA[1] | ATP | ATP | NA[1] | NA[3] | NA[1] |
| Mg | 2 | NA[2] | NA[1] | 2 | NA[2] | NA[1] | NA[3] | NA[1] |
| Water | 2 | NA[2] | NA[1] | 1 | NA[2] | NA[1] | NA[3] | NA[1] |
| **R.m.s deviation** | | | | | | | | NA[1] |
| Bond length (Å) | 0.003 | 0.003 | NA[1] | 0.005 | 0.004 | NA[1] | NA[3] | NA[1] |
| Bond angles (º) | 0.603 | 0.609 | NA[1] | 0.786 | 0.852 | NA[1] | NA[3] | NA[1] |
| **Validation** | | | | | | | | |
| Molprobity score | 1.49 | 1.43 | NA[1] | 1.53 | 1.53 | NA[1] | NA[3] | NA[1] |
| Clashscore | 5.46 | 4.60 | NA[1] | 4.55 | 5.86 | NA[1] | NA[3] | NA[1] |
| **Rotamer outliers (%)** | 0.00 | 0.00 | NA[1] | 0.00 | 0.00 | | | NA[1] |
| **Ramachandran plot** | | | | | | | | |
| Favored (%) | 96.80 | 96.80 | NA[1] | 95.65 | 96.66 | NA[1] | NA[3] | NA[1] |
| Allowed (%) | 3.20 | 3.05 | NA[1] | 4.35 | 3.34 | NA[1] | NA[3] | NA[1] |
| Outliers (%) | 0.00 | 0.15 | NA[1] | 0.00 | 0.00 | NA[1] | NA[3] | NA[1] |
| **PDB code** | 9Y4J | 9Y4L | NA[1] | 9Q6O | 9Q5W | NA[1] | NA[3] | NA[1] |
| **EMDB code** | EMD-72484 | EMD-72485 | EMD-72284 | EMD-72244 | EMD-72243 | EMD-72285 | | |

[1] Due to the low map resolution at both ends of the four-subunit filaments, models were not built for TetraMDA5/dsRNA or TetraT331I/dsRNA.

[2] $Mg^{2+}$ and water were not modeled in DiMDA5/dsRNA and DiT331I/dsRNA complex due to low resolution.

[3] Due to the low resolution of the density for AMPPNP, model was not built for MonoT331I/dsRNA/AMMPNP.

**Table S2. The sequence of in vitro transcribed RNAs that are used in luciferase assay (see Fig. 1C).**

| | |
|---|---|
| SLR30 | GGAUCGAUCGAUCGAUCGGCAUCGAUCGGCUUCGGCCGAUCGAUGCCGAUCGAUCGAUCGAUCC |
| | M-fold: ΔG = -65.40 kcal/mol |
| SLR50 | GACUACACGAAAGCUCCAAUUGUAUGCCAGGGUACAUCAUGCUGCGCAUCUUCGGAUGCGCAGCAUGAUGUACCCUGGCAUACAAUUG<br>GAGCUUUCGUGUAGUC |
| | M-fold: ΔG = -103.90 kcal/mol |
| dsRNA112 | 5'-<br>GGGCCACAUCGCAAGACAUAAAUCCAGAUGGUCAGACACAUCGGAUCGGGGCAAGGCCAAUCCCCGAUCCUGCUAUAAGUAACACACUA<br>GUCGAUCGUCUCGCGGUUCAUCC -3' |
| | 5'-<br>GGAUGAACCGCGAGACGAUCGACUAGUGUGUUACUUAUAGCAGGAUCGGGGAUUGGCCUUGCCCCGAUCCGAUGUGUCUGACCAUCU<br>GGAUUUAUGUCUUGCGAUGUGGCCC -3' |
| | M-fold: ΔG = -235.90 kcal/mol |
| dsRNA150 | 5'-<br>GGGAGAAUGUCGAAUGGGUAUUCCACAGACGAGAAUUUCCGCUAUCUCAUCUCGUGCUUCAGGGCCAGGGUGAAAAUGUACAUCCAGG<br>UGGAGCCUGUGCUGGACUACCUGACCUUUCUGCCUGCAGAGGUGAAGGAGCAGAUUUCUCCC -3' |
| | 5'-<br>GGGAGAAAUCUGCUCCUUCACCUCUGCAGGCAGAAAGGUCAGGUAGUCCAGCACAGGCUCCACCUGGAUGUACAUUUUCACCCUGGCC<br>CUGAAGCACGAGAUGAGAUAGCGGAAAUUCUCGUCUGUGGAAUACCCAUUCGACAUUCUCCC -3' |
| | M-fold: ΔG = -321.80 kcal/mol |
| dsRNA500 | 5'-<br>GGGAGAAUGUCGAAUGGGUAUUCCACAGACGAGAAUUUCCGCUAUCUCAUCUCGUGCUUCAGGGCCAGGGUGAAAAUGUACAUCCAGG<br>UGGAGCCUGUGCUGGACUACCUGACCUUUCUGCCUGCAGAGGUGAAGGAGCAGAUUCAGAGGACAGUCGCCACCUCCGGGAACAUGCA<br>GGCAGUUGAACUGCUGCUGAGCACCUUGGAGAAGGGAGUCUGGCACCUUGGUUGGACUCGGGAAUUCGUGGAGGCCCUCCGGAGAAC<br>CGGCAGCCCUCUGGCCGCCCGCUACAUGAACCCUGAGCUCACGGACUUGCCCUCUCCAUCGUUUGAGAACGCUCAUGAUGAAUAUCUC<br>CAACUGCUGAACCUCCUUCAGCCCACUCUGGUGGACAAGCUUCUAGUUAGAGACGUCUUGGAUAAGUGCAUGGAGGAGGAACUGUUGA<br>CAAUUGAAGACAGAAACCGGAUUGCUGCUGCAGAAAACAAUGGAAAUGAAUCAGGUCUCCC -3' |
| | 5'-<br>GGGAGACCUGAUUCAUUUCCAUUGUUUUCUGCAGCAGCAAUCCGGUUUCUGUCUUCAAUUGUCAACAGUUCCUCCUCCAUGCACUUAU<br>CCAAGACGUCUCUAACUAGAAGCUUGUCCACCAGAGUGGGCUGAAGGAGGUUCAGCAGUUGGAGAUAUUCAUCAUGAGCGUUCUCAAA<br>CGAUGGAGAGGGCAAGUCCGUGAGCUCAGGGUUCAUGUAGCGGGCGGCCAGAGGGCUGCCGGUUCUCCGGAGGGCCUCCACGAAUUC<br>CCGAGUCCAACCAAGGUGCCAGACUCCCUUCUCCAAGGUGCUCAGCAGCAGUUCAACUGCCUGCAUGUUCCCGGAGGUGGCGACUGUC<br>CUCUGAAUCUGCUCCUUCACCUCUGCAGGCAGAAAGGUCAGGUAGUCCAGCACAGGCUCCACCUGGAUGUACAUUUUCACCCUGGCCC<br>UGAAGCACGAGAUGAGAUAGCGGAAAUUCUCGUCUGUGGAAUACCCAUUCGACAUUCUCCC -3' |
| | M-fold: ΔG = -1109.00 kcal/mol |
